# A specialized ARGONAUTE enables trans-species RNA interference in plant immunity

**DOI:** 10.64898/2026.02.18.706620

**Authors:** Min Wang, Chao Yang, Enoch Lok Him Yuen, Xiaodong Fang, Fan Qi, Shuang Yang, Benjamin L Koch, Kun Li, Yingnan Hou, Taerin Oh, Bozeng Tang, Li Feng, Xiuren Zhang, Tolga Osman Bozkurt, Xiaoqi Feng, Wenbo Ma

## Abstract

Trans-species RNA interference (tsRNAi), in which plants produce small RNAs (sRNAs) to silence target genes in pathogens, has emerged as a promising strategy for disease control. However, whether tsRNAi constitutes an endogenous, regulated immune response remains unclear. Here, we show that ARGONAUTE10 (AGO10) plays a critical role in pathogen-induced tsRNAi. Loss of AGO10 in Arabidopsis abolished pathogen gene silencing during infection, leading to hypersusceptibility to oomycete and fungal pathogens. Importantly, AGO10 rapidly responds to pathogen infection through increased protein accumulation and re-location into discrete cytoplasmic condensates, thus promoting the production of trans-species sRNAs at the pathogen infection sites. This immune responsiveness relies on the N terminal intrinsically disordered region (IDR) of AGO10, which is responsible for sensing and responding to immune activation. Specific features in the IDR partitions AGO10 into two deeply diverged subgroups, AGO10a and AGO10b, with the immune responsiveness and defense function evolutionarily conserved in AGO10a but not AGO10b. Together, these findings establish tsRNAi as a bona fide, evolutionarily conserved immune response and position AGO10 as a signal-responsive hub linking pathogen perception to tsRNAi-based defense.

## Introduction

Small RNA (sRNA)-guided gene silencing regulates diverse biological processes in eukaryotes, including development, stress responses, and genome integrity maintenance(1, 2). Beyond endogenous gene regulation, growing evidence supports that plant-produced sRNAs can act as mobile silencing agents to suppress pathogen gene expression(3). This phenomenon of trans-species RNAi (tsRNAi) underlies host-induced gene silencing (HIGS), a strategy in which plants are engineered to produce double-stranded RNAs (dsRNAs) or artificial sRNAs designed to silence selected pathogen targets(4, 5). Despite its promise in crop protection, molecular mechanisms governing tsRNAi remain poorly understood, limiting an effective implementation of HIGS in agriculture.

Plant sRNAs, usually ranging from 20 to 24 nucleotides (nt) in size, are classified into microRNAs (miRNAs) and small interfering RNAs (siRNAs) according to their distinct biogenetic pathways(6, 7). Both miRNAs and siRNAs can be loaded onto ARGONAUTE (AGO) proteins to guide sequence-specific target gene silencing(8), but many sRNAs with demonstrated tsRNAi activity during pathogen infection are 21-nt secondary siRNAs(3). Biogenesis of secondary siRNAs is initiated by miRNA-guided and AGO-dependent “slicing” of a target poly(A) transcript. The cleaved product then serves as the template for the synthesis of dsRNA by RNA-dependent RNA polymerase 6 (RDR6). The dsRNA is further processed into 21-nt increments of siRNAs, which are also called phased siRNAs or phasiRNAs(7, 9). In the model plant *Arabidopsis thaliana*, mutants defective in secondary siRNA production are hypersusceptible to oomycete and fungal pathogens(10-12). Pathogens, in turn, have evolved virulence proteins that suppress the secondary siRNA pathway in their hosts(11, 13, 14). These findings implicate secondary siRNAs as major executors of tsRNAi. However, how these siRNAs, and tsRNAi in general, are regulated during pathogen infection is unknown. Therefore, whether tsRNAi is a bona fide immune response remains unclear.

## Results

### AGO10 is required for tsRNAi during pathogen infection

AGOs are evolutionarily conserved endonucleases and central players in sRNA-guided gene silencing(8). In plants, AGOs with “slicer” activity are also required to initiate secondary siRNA production, making them promising candidates in regulating tsRNAi. *A. thaliana* encodes ten AGOs. We conducted a screen in the Col-0 background and revealed three mutants, *ago1-45* (a weak allele of AGO1 as null mutations cause embryonic lethality), *ago7*, and *ago10-1*, that exhibited hypersusceptibility to the oomycete pathogen *Phytophthora capsici* (Fig. 1a and Fig. S1a). Among them, AGO10 was not known to contribute to defense. We confirmed the *ago10* phenotype by testing an independent mutant *ago10-2*, which was also hypersusceptible to *P. capsici* (Fig. 1b and Fig. S1b). In addition, both *ago10* mutants showed increased susceptibility to the fungal pathogen *Colletotrichum higginsianum* (Fig. S1c,d), indicating a broad contribution of AGO10 to pathogen resistance.

**Figure 1.**
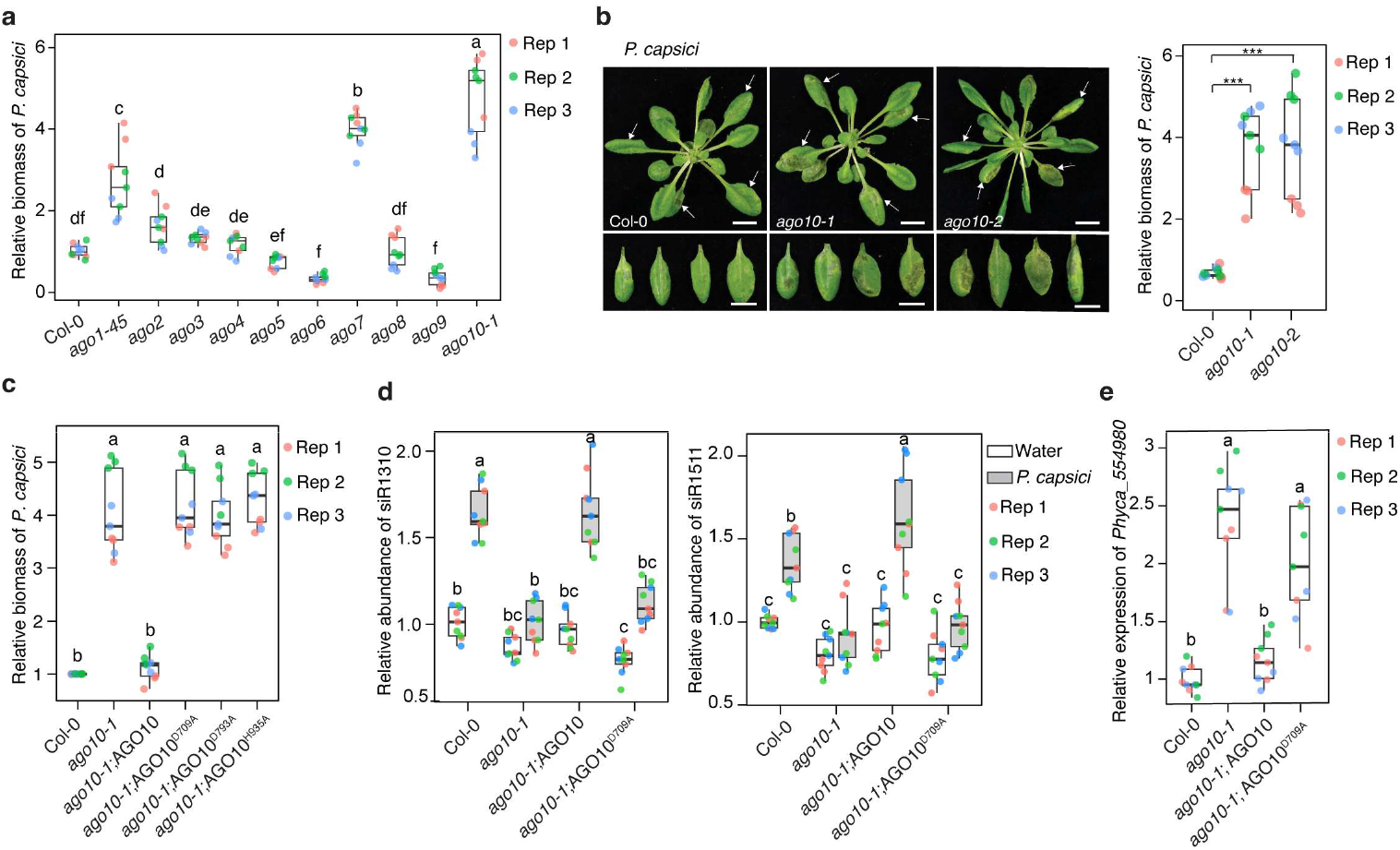
AGO10 contributes to plant defense through trans-species gene silencing. **a**, Mutation in *AGO10* led to hypersusceptibility to *Phytophthora capsici*. Four-week-old plants of wildtype (Col-0) and ten *ago* mutants in *A. thaliana* were inoculated with zoospore suspension. Pathogen biomass was determined at 3 days post-inoculation (dpi) by qPCR in each genotype. Relative biomass was calculated by comparing to the biomass in wildtype plants. **b**, Enhanced susceptibility of *ago10* mutants to *P. capsici*, represented by disease symptoms (left) and pathogen biomass (right). Arrowheads indicate the inoculated leaves. Scale bars, 2.0 cm. **c**, Catalytic mutants (AGO10^D709A^, AGO10^D793A^, and AGO10^H935A^) of AGO10 were unable to rescue the hypersusceptibility phenotype of *ago10-1* mutant. **d**, AGO10 is required for increased siRNA accumulation during *P. capsici* infection. Two phasiRNAs, siR1310 and siR1511, were evaluated using stemloop-PCR with *AtACTIN* as the internal reference. **e**, AGO10 is required for the silencing of the *P. capsici* gene *Phyca_554980* during infection. Transcript abundance of *Phyca_554980* from inoculated tissue of *A. thaliana* was determined by qRT-PCR with *Pc76RT* as the internal reference. In **a**–**e**, data from three biological replicates are presented; different letters indicate statistically significant differences (*p* < 0.05) determined by one-way ANOVA with Tukey’s multiple comparisons test. In **b**, *p* values were calculated using two-tailed Student’s *t* test (\*\*\**p* < 0.001).

AGO10 contains a conserved Asp-Asp-His (DDH) catalytic triad for RNA cleavage(15, 16). Previous studies show that AGO10 antagonizes the function of AGO1 and protects the targets of miR165/166 from being silenced in meristem and vasculature tissues of some *A. thaliana* ecotypes(15, 17-20). For example, in Ler background, loss of *AGO10* led to the “pinhead” phenotype, which can be fully rescued by catalytic-dead mutants, consistent with this unique regulatory mechanism(15). In contrast, none of the catalytic mutants were able to restore resistance to *P. capsici* (Fig. 1c and Fig. S1e,f). It is worth noting that there are no visible developmental defects in the *ago10* mutants in Col-0 background, which is what we used for all our experiments. These results indicate that AGO10 functions through a catalytic activity-dependent mechanism in plant immunity, contrasting its known function in development.

We further examined the role of AGO10 in pathogen gene silencing. Secondary siRNAs derived from *PPR* and *TAS* transcripts in *A. thaliana* have been implicated in tsRNAi during *P. capsici* infection(11, 21). We quantified two representative siRNAs, siR1310, derived from *PPR* transcripts, and siR1511, derived from *TAS2*, in *A. thaliana* leaf tissues following *P. capsici* infection. Both siRNAs showed increased accumulation in wildtype Col-0 plants after pathogen inoculation; however, these inductions were abolished in *ago10-1* (Fig. 1d). siR1310 targets the *P. capsici* U2 splicing factor gene *Phyca_554980* for silencing during infection and this tsRNAi contributes to defense(11). Consistent with the reduced siR1310 levels, *Phyca_554980* transcript level was significantly higher in *P. capsici* infected *ago10-1* than Col-0 (Fig. 1e). In both experiments, the catalytic mutant AGO10^D709A^ was unable to complement the *ago10-1* mutant phenotypes (Fig. 1d,e). These results demonstrate that AGO10 is required for infection-induced accumulation of trans-species siRNAs and the silencing of pathogen genes.

### AGO10 undergoes rapid and localized activation in response to pathogen infection

The increased accumulation of trans-species siRNAs after pathogen infection prompted us to examine whether AGO10 displays immune-responsive regulation. RT-qPCR results demonstrated that *AGO10* transcript levels remained unchanged after *P. capsici* inoculation (Fig. S2a). Consistently, single-cell transcriptome data(22) revealed no transcriptional changes across all cell types during the infection of *C. higginsianum* (Fig. S2b), indicating that *AGO10* is not transcriptionally activated after pathogen infection.

We next monitored AGO10 protein dynamics in *A. thaliana* leaf tissue inoculated with *P. capsici* zoospores using transgenic plants expressing *p35S::Flag-4Myc-AGO10*. In mock (water)-treated plants, AGO10 proteins accumulated at a relatively low level; however, a significant increase was observed as early as 1 hour post inoculation (hpi) and this induction was maintained till at least 9 hpi (Fig. 2a). This drastic increase of AGO10 proteins was also detected after treatment by pathogen-associated molecular patterns (PAMPs) including the bacterial flagellin epitope flg22 and the fungal cell wall component chitin (Fig. 2b and Fig. S2c,d). Similarly, a strong induction at 30 minutes after flg22 treatment was also observed when *AGO10* was expressed under its native promoter (Fig. 2c). These results suggest that AGO10 is induced at the post-transcriptional level upon detection of pathogen invasion, representing an immune response. In comparison, AGO1 protein level remained unchanged during *P. capsici* infection (Fig. S2e), indicating a specific role of AGO10 in plant immunity.

**Figure 2.**
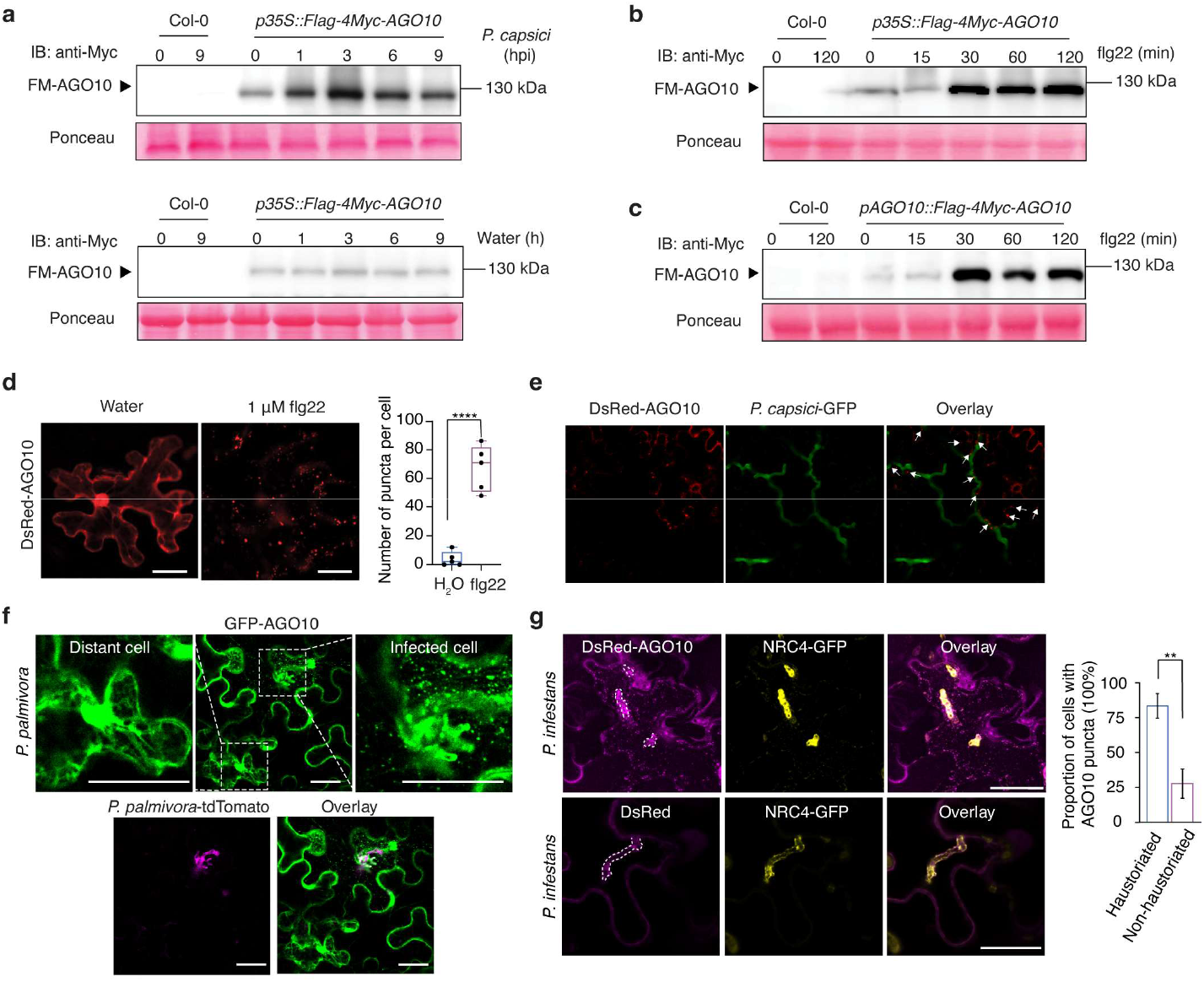
AGO10 is responsive to pathogen infection. **a**, Increased accumulation of AGO10 proteins in *A. thaliana* following *P. capsici* inoculation. **b,c**, Increased accumulation of AGO10 proteins in *A. thaliana* treated with 1 μM flg22. In **a**–**c**, ten-day-old seedlings of transgenic plants expressing *p35S::Flag-4Myc-AGO10* or *pAGO10::Flag-4Myc-AGO10* were examined at the indicated timepoint by immunoblotting using anti-Myc antibody. Ponceau S staining served as the loading control. **d**, Con-focal images showing cytoplasmic puncta formed by AGO10 in response to flg22 treatment. DsRed-AGO10 was expressed in *Nicotiana benthamiana*. Forty-eight hours post Agroinfiltration, the leaves were treated with flg22 and images were taken after 10 minutes. Scale bars, 25 μm. *p* values were calculated using two-tailed Student’s *t* test (\*\*\*\**p* < 0.0001, \*\**p* < 0.01). **e**, Association of AGO10 puncta (indicated by arrowheads) with *P. capsici* hyphae during infection. *N. benthamiana* expressing DsRed-AGO10 was inoculated with zoospores, and the images were taken at 12 hpi. Scale bars, 25 μm. **f**, AGO10 puncta were observed in plant cells at the infection site of *Phytophthora palmivora. N. benthamiana* expressing GFP-AGO10 was inoculated with zoospores and the images were taken at 12 hpi; Magnified views of boxed regions showing a cell in direct contact with *Phytophthora* hyphae (right) and a more distant, uninfected cell (left). Scale bars, 25 μm. **g**, AGO10 puncta were formed in haustoriated cells during the infection of *Phytophthora infestans*. NRC4-GFP was co-expressed with DsRed-AGO10 and used to mark the extrahaustorial membrane. Images were taken at 3 dpi. Scale bars, 25 μm. Statistical significance was assessed using Fisher’s exact test (n = 18 images per condition, \*\**p* < 0.01).

To further explore the immune responsiveness of AGO10, we investigated its subcellular localization after immune activation. DsRed-AGO10 transiently expressed in *Nicotiana benthamiana* showed a diffused cytoplasm-nuclear distribution under untreated conditions. Strikingly, flg22 treatment induced rapid re-localization of AGO10, forming distinct cytoplasmic puncta within 10 minutes (Fig. 2d). This change in protein localization was accompanied by reduced fluorescent signals in the nuclei. Similar cytoplasmic puncta formation was also observed following the infection of various pathogens, including *P. capsici* (Fig. 2e and Fig. S2f), *Phytophthora palmivora* (Fig. 2f and Fig. S2g), *Phytophthora infestans* (Fig. 2g), and *C. higginsianum* (Fig. S2h). Through these experiments, we observed that the plant cells in which AGO10 formed cytoplasmic puncta were often in direct contact to the invasive hyphae, while the more distant cells retained the diffused cytoplasm-nuclear distribution, indicating that AGO10 re-localization is a localized response at infection sites. We further strengthened this observation by examining co-appearance of AGO10 puncta with haustoria, which are specialized infection structures formed by *Phytophthora* and other filamentous pathogens. AGO10 was co-expressed in *N. benthamiana* with the solanaceous NUCLEOTIDE-BINDING LEUCINE-RICH REPEAT (NLR) protein NLR REQUIRED FOR CELL DEATH 4 (NRC4), which accumulates at the extrahaustorial membrane that encases haustoria formed by *P. infestans*(23). Using NRC4 as a marker, we found that 93.75% of imaged haustoria (n=45) had AGO10 puncta. Furthermore, AGO10 puncta were significantly enriched in infected haustoriated regions, being detected in 83.3% haustoriated cells, compared to only 27.7% of uninfected cells in the same leaf had AGO10 puncta (Fig. 2g). Together, these results demonstrate a spatial-temporal dynamic of AGO10 proteins during pathogen infection, indicating an immune-activated functional switch.

### AGO10 is responsible for immune-induced trans-species siRNA production

The cytoplasmic puncta formed by AGO10 following immune activation are reminiscent of siRNA bodies, which are membraneless cytoplasmic organelles associated with secondary siRNA biogenesis(24). siRNA bodies are typically characterized by the presence of SUPPRESSOR OF GENE SILENCING 3 (SGS3), a key scaffolding protein in the RDR6-dependent siRNA pathway(24). AGO1 and AGO7, which have known function in the initial step of secondary siRNA biogenesis, were found to associate with SGS3 in the siRNA bodies(25). By co-expressing AGO10 with SGS3 in *N. benthamiana*, we observed a strong co-localization in cytoplasmic puncta (Fig. 3a). AGO10 also associated with SGS3 in plant cells, as demonstrated by co-immunoprecipitation (Fig. 3b) and bimolecular fluorescence complementation (BiFC) (Fig. S3a). These results indicate that the cytoplasmic puncta formed by AGO10 are likely siRNA bodies.

**Figure 3.**
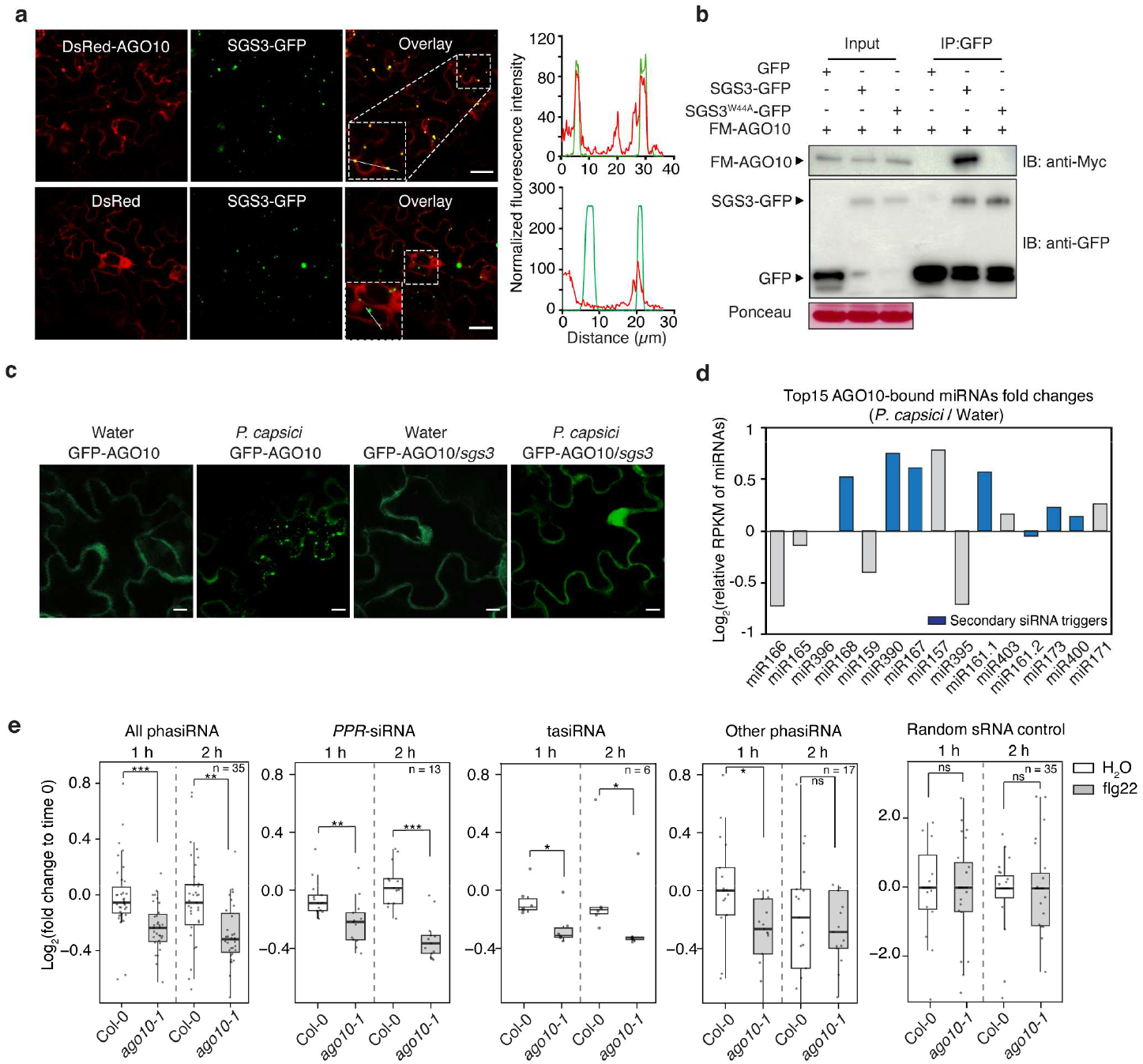
AGO10 is required for immune-induced siRNA production. **a**, AGO10 co-localizes with the siRNA body marker SGS3 in cytoplasmic foci. AGO10 and SGS3 were co-expressed in *N. benthamiana*. Line-scan intensity profiles were analysed using ImageJ. Scale bars, 25 μm. **b**, AGO10 interacts with SGS3 through a predicted GW motif in SGS3. Wildtype SGS3, but not the mutant SGS3^W44A^, was co-precipitated with AGO10 when co-expressed in *N. benthamiana*. **c**, Puncta formation of AGO10 depends on SGS3. Transgenic plants expressing GFP-AGO10 in wildtype (Col-0) or *sgs3* mutant background were inoculated with *P. capsici* and examined for AGO10 localization. Scale bars, 25 μm. **d**, AGO10 binds to miRNA triggers of secondary siRNA production. AGO10 proteins were immunoprecipitated from four-week-old *A. thaliana* with or without *P. capsici* infection 8 hpi for sRNA-seq analysis. miRNAs are ordered based on their abundance from high (left) to low (right). Blue bars indicate miRNAs capable of triggering secondary siRNA biogenesis. Data represent averages from three biological replicates. **e**, Reduced accumulation of phasiRNAs in *ago10* compared to wildtype (Col-0) plants after flg22 treatment. Ten-day-old *A. thaliana* seedlings were treated with 1 μM flg22. Samples were collected at 0, 1 and 2 hours for sRNA-seq. Relative RPKM of sRNAs from all 35 loci that produced siRNAs (including 13 *PPR* and 6 *TAS*)(31) were analyzed. Thirty-five non-PHAS 21-nt sRNA clusters were randomly selected from ShortStack(32)-identified clusters as negative controls, excluding annotated PHAS. Pairwise comparisons were performed using unpaired Wilcoxon rank-sum tests with Benjamini–Hochberg correction for multiple testing. \*\*\**p* < 0.001; \*\**p* < 0.01, \**p* < 0.05; ns, not significant.

Using AlphaFold-Multimer(26), we generated a structural model of the AGO10-SGS3 protein complex, which revealed a high-confidence interaction interface between a putative GW motif(27) from SGS3 and a binding pocket in AGO10 (Fig. S3b). Six amino acids in AGO10 were predicted to directly interact with W44 in the predicted SGS3 GW motif. Interestingly, the predicted AGO10-SGS3 interaction interface resembles the well-characterized interaction between human hAGO1 and hGW182(28) (Fig. S3c). To validate this model, we generated a mutant SGS3^W44A^. Using Co-immunoprecipitation, we found that SGS3^W44A^ lost the ability to interact with AGO10 (Fig. 3b). Furthermore, AGO10 could no longer form cytoplasmic puncta when co-expressed with SGS3^W44A^ (Fig. S3d), indicating that SGS3 interaction is required for the re-localization of AGO10 to siRNA bodies. Consistent with this hypothesis, GFP-AGO10 remained the diffused cytoplasm-nuclear distribution in *A. thaliana sgs3* mutant after *P. capsici* infection (Fig. 3c), although the AGO10 protein level was still induced (Fig. S3e). These results suggest that AGO10 is recruited to siRNA bodies after immune activation through a direct association with SGS3.

Secondary siRNA biosynthesis is initiated by specific interactions of an AGO with trigger miRNAs(29, 30). We therefore analyzed miRNAs associated with AGO10 after *P. capsici* infection in *A. thaliana*. AGO10 was immunoprecipitated from leaf tissues of transgenic plants expressing Flag-4Myc-AGO10 at 8 hpi and the associated miRNAs were determined by sRNA-seq (Fig. S4a,b). Consistent with previous reports(15), AGO10-bound sRNAs showed a predominant enrichment of 5’ terminal uridine (U) (Fig. S4c). We found several miRNAs, such as miR165/166, miR168 and miR173, that are known to associate with AGO10 in meristem(15, 17). Importantly, we observed that miRNAs known to trigger secondary siRNA production exhibited increased loading into AGO10 after *P. capsici* infection (Fig. 3d). Of particular interest is miR161 and miR173, which trigger the production of trans-species siRNAs derived from *PPR* and *TAS* transcripts. This infection-induced shift in AGO10-associated miRNAs, together with re-localization in the siRNA bodies and the requirement of catalytic activity for its function in tsRNAi, suggests that AGO10 may directly participate in the biogenesis of trans-species siRNAs during pathogen infection.

To directly examine the effect of AGO10 on siRNA accumulation, we performed sRNA-seq using wildtype and *ago10-1* mutant plants with flg22 treatment (Fig. S5a–c). Leaf tissues were collected at 1 and 2 hours post treatment and the abundance of siRNAs were quantified relative to time 0. Our results revealed significant reductions of secondary siRNAs in *ago10-1*, especially phasiRNAs derived from *PPR* and *TAS* transcripts, at both time points (Fig. 3e). In contrast, randomly selected sRNA-producing loci did not show difference. These results were further confirmed by quantifying siR1310 using qRT-PCR. In wildtype plants, flg22 treatment led to an increase in siR1310 but this induction was abolished in *ago10-1* (Fig. S5d). Furthermore, the induced accumulation of siR1310 could be complemented by wildtype but not catalytic mutant of AGO10 (Fig. S5d). Together, these results indicate that AGO10 plays a key role in immune-induced production of secondary siRNAs.

### An N-terminal IDR is required for immune-responsiveness and defense function of AGO10

SGS3-containing siRNA bodies have recently been shown to assemble through liquid–liquid phase separation (LLPS)(25). We therefore examined biophysical properties of the immune-induced AGO10 puncta and tested whether they also represent dynamic biomolecular condensates. DsRed-AGO10 was expressed in *N. benthamiana*, and the leaves were subsequently inoculated with *P. capsici* before analysis by fluorescence recovery after photobleaching (FRAP). AGO10 fluorescence signal rapidly recovered following photobleaching with approximately 90% of the initial signal restored within 3 minutes (Fig. 4a,b). This observation indicates highly fluid molecular exchange that is characteristic of liquid-like structures. Time-lapse confocal microscopy captured spontaneous fusion of AGO10 condensates upon contact (Fig. 4c), another hallmark of liquid-like droplets. These observations suggest that LLPS underlies the immune-induced AGO10 puncta formation.

**Figure 4.**
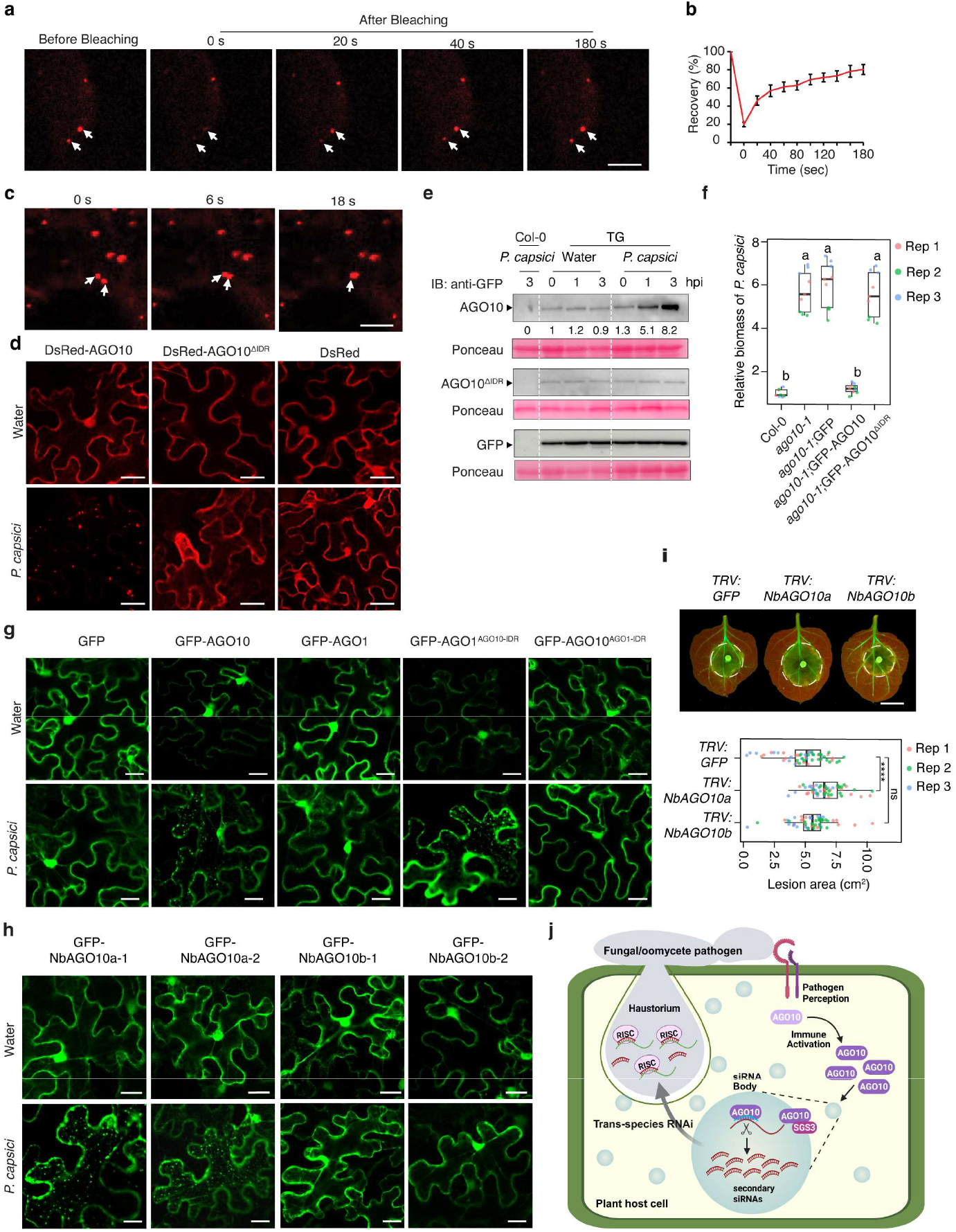
An N-terminal IDR is essential for immune-responsiveness and defense function of AGO10. **a,b** AGO10 puncta show liquid-like properties. DsRed-AGO10 was expressed in *N. benthamiana* and inoculated with *P. capsici* before analzyed by fluorescence recovery after photobleaching (FRAP). Scale bar, 5 μm. In **b**, Data represent mean ± SEM from four independent experiments. **c**, Time-lapse fluorescence microscopy detected fusion events (indicated by arrowheads) between AGO10 puncta. Scale bar, 5 μm. **d**, The N-terminal IDR is required for AGO10 puncta formation. Scale bars, 25 μm. **e**, The IDR is required for immune-induced accumulation of AGO10 proteins. Ten-day-old *A. thaliana* seedlings were inoculated with *P. capsici* and AGO10 proteins were monitored by immunoblotting using an anti-GFP antibody. Ponceau S staining was used as the loading control. TG, transgenic plants expressing GFP-AGO10, GFP-AGO10^ΔIDR^ or GFP in *ago10-1* mutant. **f**, GFP-AGO10^ΔIDR^ was unable to complement the hypersusceptibility phenotype of *ago10-1* mutant. Four-+old *A. thaliana* plants were inoculated with *P. capsici* and the pathogen biomass was determined at 3 dpi. Different letters indicate significant differences (*p* < 0.05, one-way ANOVA with Tukey’s multiple comparisons). **g**, IDR of AGO10 is sufficient to drive infection-induced puncta formation. Scale bars, 25 μm. Con-focal images show subcellular localizations of AGO1, AGO10 and the chimeras when expressed in *N. benthamiana*. **h**, NbAGO10a, but not NbAGO10b, form cytoplasmic puncta after transient expression in *N. benthamiana* and *P. capsici* infection. Scale bars, 25 μm. **i**, NbAGO10a, but not NbAGO10b, contribute to defense. Disease phenotypes were assessed in *N. benthamiana* plants silenced for *NbAGO10a* or *NbAGO10b* using virus-induced gene silencing (VIGS). Representative images showing lesions were captured at 2 dpi. Scale bar, 2 cm. Lesion areas were quantified using ImageJ. Data were analysed by two-tailed Student’s *t*-test (\*\*\*\**p* < 0.0001, ns: not significant, n ≥ 15 leaves per genotype per experiment). **j**, A proposed model illustrating AGO10 as a critical component of trans-species RNAi-based immunity. Pathogen perception and immune activation led to a rapid accumulation of AGO10 proteins, which are subsequently recruited to siRNA bodies, in which AGO10 promotes secondary sRNA biogenesis at the pathogen infection sites. Immune responsiveness of AGO10 depends on an N-terminal IDR and the re-localization to siRNA bodies also requires interaction with SGS3. AGO10-dependent siRNAs elevate pathogen resistance through tsRNAi. RISC: RNA-induced silencing complex. The figure was created in Biorender https://BioRender.com/eg8co3h.

Intrinsically disordered regions (IDRs) frequently drive LLPS(33).AGOs are known to contain IDRs, especially at the N terminus(34, 35). We predicted an IDR domain within the N-terminal 1-125 amino acids of AGO10 (Fig. S6a) and assessed the contribution of this region to AGO10 puncta formation. When expressed in *N. benthamiana*, a mutant lacking the N-terminal IDR (AGO10^ΔIDR^) failed to form condensates after *P. capsici* inoculation (Fig. 4d and Fig. S6b). Furthermore, GFP-AGO10^ΔIDR^ expressed in *A. thaliana ago10-1* mutant background remained diffuse in the cytosol after *P. capsici* infection (Fig. S6c,d). Deleting the N-terminal IDR also abolished the induced protein accumulation of AGO10 (Fig. 4e). Consistent with these results, AGO10^ΔIDR^ was unable to rescue the hypersusceptibility phenotype of *ago10-1* (Fig. 4f and Fig. S6e,f). These results indicate that the immune-responsiveness of AGO10 requires the N-terminal IDR and potentially an LLPS-dependent process.

Eukaryotic AGOs share a highly conserved architecture, which generally consists of four functional domains: N-terminal (N) domain, PAZ, MID and PIWI(8). The AGO10 IDR is located upstream of the N domain, which is the region with the highest sequence divergence across the ten *A. thaliana* AGOs (Fig. S7a). The closest homolog of AGO10 is AGO1, which also contains an IDR at its N terminus but barely shares any sequence similarity with AGO10 IDR (Fig. S7b,c). AGO1 exhibits a diffused cytoplasm-nuclear distribution but could form condensates after heat treatments(34). However, unlike AGO10, the localization of AGO1 remained the same after *P. capsici* inoculation (Fig. 4g). We constructed chimeras by swapping the N-terminal regions of AGO1 and AGO10. Remarkably, AGO1 containing the AGO10 IDR (AGO1^AGO10-IDR^) acquired the ability to form cytoplasmic puncta in response to *P. capsici* infection, similar to wildtype AGO10, whereas AGO10 containing the AGO1 IDR (AGO10^AGO1-IDR^) lost this response (Fig. 4g). These results demonstrate that the N-terminal IDR of AGO10 is necessary and sufficient for sensing and responding to immune activation. Interestingly, introducing the AGO1^AGO10-IDR^ chimera into *ago10-1* did not rescue the hypersusceptible phenotype (Fig. S7d–g), indicating that additional feature(s) of AGO10 is required for its defense function.

### AGO10 has an evolutionarily conserved function in immunity

AGOs in angiosperms are classified into three major phylogenetic clades: AGO1/5/10, AGO2/3/7, and AGO4/6/8/9(36). Recent analyses across the green lineages suggest that AGO10 represents the ancestor-like form in the AGO1/5/10 clade. This AGO10-like ancestor subsequently diverged into two subclades, AGO10a and AGO10b. While most angiosperms encode both AGO10a and AGO10b, Brassicaceae species, including *A. thaliana*, lost AGO10b and only encodes AGO10a(36) (Fig. S8a). Considering the importance of the N-terminal IDR in the defense function of *A. thaliana* AGO10, we analyzed members of both AGO10a and AGO10b subclades from representative plant species with a focus on their N-terminal sequences. This analysis revealed that AGO10a proteins share a proline-rich domain (PRD), which is absent in AGO10b proteins (Fig. S8b–d). The only exception is rice, which encodes a single AGO10 that belongs to the AGO10b subclade but contains a PRD. Furthermore, the liverwort *Marchantia polymorpha* encodes only one AGO in AGO1/5/10 clade and this AGO has a longer IDR that contains multiple PRDs. Proline-rich regions are known to facilitate LLPS(33), we therefore tested whether other AGO10a members can also form immune-induced condensation and contribute to defense. For this purpose, we examined the AGO10 orthologs of *Nicotiana benthamiana*, which encodes two AGO10a and two AGO10b. The NbAGO10 orthologs were expressed in *N. benthamiana* and their subcellular localizations were monitored after *P. capsici* inoculation. Without infection, all NbAGO10 proteins showed a diffused cytoplasm-nuclear distribution. After infection, the two NbAGO10a proteins underwent re-localization and formed cytoplasmic puncta, while the NbAGO10b proteins remained unchanged (Fig. 4h). Furthermore, silencing of *NbAGO10a*, but not *NbAGO10b*, led to hypersusceptibility when inoculated with *P. capsici* (Fig. 4i and Fig. S8e,f). Together, these results indicate that the function in plant immunity may be evolutionarily conserved in AGO10a subclade and suggest that divergence in the N-terminal IDR may underline functional specialization among AGO10 paralogs.

## Discussion

A defining feature of immune systems is their ability to remain quiescent during normal growth yet rapidly activate upon pathogen detection. Although tsRNAi have been implicated in plant–pathogen interactions, whether it is dynamically regulated as an active defense mechanism remains unclear. Here we discovered a specialized ARGONAUTE as a molecular link between pathogen perception and tsRNAi-based plant immunity, establishing tsRNAi as a well-controlled defense response. Our finding supports a model in which plants maintain a basal level of secondary siRNAs that serve as a surveillance system. During pathogen infection, AGO10 rapidly accumulates in immune-activated cells and is recruited to SGS3-dependent siRNA bodies, promoting secondary siRNA production and pathogen gene silencing at the infection sites (Fig. 4j). The localized accumulation of antimicrobial siRNAs intensifies defenses against pathogens while minimizing unintended effects on plant growth and beneficial microbiomes. This mechanistic insight provides critical guidance in deploying tsRNAi-based immunity for crop protection.

A striking feature of AGO10 immune responsiveness is its rapid protein accumulation and re-localization into siRNA bodies, both depending on an N-terminal IDR. As such, AGO10’s IDR serves as a sensor for immune activation, potentially through post-translational modification, protein-protein interaction(s), and/or LLPS. A key component of siRNA bodies is SGS3, which is a scaffolding protein that assembles secondary siRNA production machinery including AGOs and RDRs(25, 37). It remains to be determined how AGO10’s IDR mediates its activation upon pathogen invasion and enables its recruitment to the siRNA bodies in coordination with SGS3 interaction.

As a functional endonuclease, the previously known function of AGO10 in regulating development does not require its enzymatic activity(15, 19, 20). This work suggests that the maintenance of AGO10 as an active enzyme is attributed to its new role in immunity, which also supports that this function in plant defense is conserved in AGO10a orthologs. AGO10 can load multiple miRNAs capable of triggering secondary siRNA production. It was reported that binding selectivity of AGO10 to its physiological substrates can be shaped by metabolites and chaperone proteins *in vitro*(38). Pathogen infection may create a specific cellular environment that facilitates routing of the miRNA triggers into AGO10 for siRNA production. Further analysis is required to understand how AGO10 is engaged in the regulation of development and immunity through distinct molecular mechanisms.

## Methods

### Plant materials and growth conditions

*Arabidopsis thaliana* ecotype Columbia (Col-0, wild-type), mutants, and transgenic plants were grown in a controlled environment chamber at 20°C under either long-day (16-hour light/8-hour dark) or short-day (10-hour light/14-hour dark) conditions. *Nicotiana benthamiana* wild-type plants were cultivated in a controlled environment chamber under a 16-hour light/8-hour dark photoperiod at 22°C.

### Pathogens and growth conditions

*Phytophthora capsici* (isolate LT263) was routinely cultured on V8 medium in a growth chamber at 25°C in darkness based on previous study(44). For zoospore production, mycelial plates were sectioned into plugs after 2 days of growth. The plugs were washed six times with sterile water for 30 minutes per wash, then incubated in sterile water at 25°C for 48 hours in darkness. Zoospore release was induced by exposing mycelia to 4°C for 40 minutes followed by room temperature for 20 minutes. The zoospore suspension was adjusted to 5 x 10^5^ zoospores/for inoculation.

*Colletotrichum higginsianum* (isolate IMI 349061) was cultured on Potato Dextrose Agar (PDA) plates at 25°C under a 12-hour light/12-hour dark cycle as previously described(45). For conidial production, one-week-old fungal cultures were flooded with room temperature water containing 0.2% gelatin. Conidia were harvested by gently scraping the culture surface with a sterile spreader. The conidial suspension was adjusted to 5 x 10^5^ conidia/mL for inoculation.

*Phytophthora palmivora* ARI-tdTomato (P3914) was maintained and zoospores were collected as previously described(46). For zoospore production, 7- to 10-day-old cultures on V8 juice agar plates were flooded with sterile distilled water at 4°C for 30 minutes, followed by an additional 30-minute incubation at room temperature (22-25°C) to trigger zoospore release. The zoospore suspension was adjusted to 5 x 10^5^ zoospores/mL for inoculation.

*Phytophthora infestans* strains (isolate 88069) were cultured on rye sucrose agar (RSA) medium in darkness at 18°C for 10-15 days prior to zoospore harvest. For zoospore production, cold water (4°C) was added to the cultures, which were then incubated in darkness at 4°C for 2 hours. For infection assays, 10 µL droplets of zoospore suspension (5 x 10^5^ zoospores/mL) were applied to the abaxial surface of agroinfiltrated leaves.

### Plasmid construction and plant transformation

Genes of interest were cloned into the expression vector *pICSL86977OD* using the In-Fusion™ HD Cloning System (Takara Bio, USA) for stable expression in *A. thaliana* and transient expression in *N. benthamiana*. DsRed and GFP coding sequences were amplified from vectors *pGDR* and *pICSL50008*, respectively, and fused to the N-terminus of AGO10 coding sequences. The Gibson Assembly® Cloning System was used to introduce SGS3 and fluorescent protein tags into the destination vector *pICSL86977OD*.

### *A. thaliana* and *N. benthamiana* infection assays

Four-week-old *A. thaliana* plants grown under short-day conditions were used for pathogenicity assays. For *P. capsici* infection, the zoospore suspension was applied to the abaxial surface of leaves using a paintbrush. Sterile water was applied as the mock control. The inoculated plants were covered with transparent lids to maintain high humidity and incubated in darkness at room temperature for 24 hours before transferring to short-day growth conditions. Disease symptoms were assessed at 3 days post-inoculation, and leaf tissue samples were harvested for pathogen biomass quantification. The relative biomass of *P. capsici* was determined by quantitative reverse transcription PCR (RT-qPCR) using *P. capsici*-specific primers, with *Arabidopsis RUB4* serving as the internal reference gene. All primer sequences are listed in Supplementary Table 1. For *C. higginsianum infection*, a 10 μL conidial suspension was spotted onto the adaxial surface of leaves of *A. thaliana*. The inoculated plants were covered with transparent lids to maintain high humidity and then incubated for seven days under short-day conditions before water-soaked lesions were assessed and photographed. Disease severity, represented by lesion area, was quantified using ImageJ.

For *N. benthamiana* infection assays, either pathogen zoospore suspensions or mycelial plugs (0.5 cm diameter) were applied to the abaxial surface of detached leaves from four-week-old plants. The inoculated leaves were maintained on moistened filter paper in darkness at 25°C. Lesion areas were measured at 2 dpi, photographed, and quantified using ImageJ software or the leaves were used for microscopy imaging.

### AGO10 complex immunoprecipitation and small RNA purification

Four-week-old *A. thaliana* plants expressing *p35S::Flag-4Myc-AGO10* were inoculated with *P. capsici*, and the infected leaves were harvested for AGO10 complex isolation 8 hpi. Tissues from water-treated plants served as the mock control. 5 g of plant material was ground to a fine powder in liquid nitrogen, and total proteins were extracted using two volumes of extraction buffer (150 mM Tris-HCl, pH=7.5, 150 mM NaCl, 1 mM MgCl_2_, 1 mM EDTA, 10% Glycerol, 5 mM DTT, 0.1% NP40, 25 μM MG132, and EDTA-free protease inhibitor cocktail). The extracts were incubated with 50 μL of pre-equilibrated anti-Flag M2 affinity beads at 4°C for 1.5 hours. After three washes, the beads were transferred to 2 mL tubes, centrifuged to remove the supernatant, and then treated with 100 μL proteinase K at 25°C for 15 mins in a heat block with shaking at 700 rpm. Finally, 700 μL QIAzol lysis reagent from the miRNeasy Micro Kit was added to the tubes. The bead-containing lysate could be stored at −80°C or used directly for small RNA extraction.

RNA from AGO10 complexes was extracted using the miRNeasy Micro Kit (QIAGEN), following the manufacturer’s instructions, and eluted from the columns using 15 μL RNase-free water.

### Small RNA library sequencing and data analysis

Small RNA library sequencing and data analysis was performed as previously described with minor modifications(47). sRNA libraries were prepared using RealSeq-Biofluids NGS Library Preparation Kit. Size-selection and purification was then performed using Novex TBE 6% polyacrylamide gels to enrich for sRNA fragments ranging from 20 to 30 nt in length. The concentration and size distribution of the purified sRNA libraries were analysed using high sensitivity DNA chips.

The sRNA-seq data was analysed as previously described(48). Adapter sequences were predicted using DNApi(49) and subsequently trimmed using Cutadapt v1.16(50). The reads were then mapped to the *Arabidopsis thaliana* reference genome (Araport11)(51) using BowTie v1.2.1.1(52) with stringent parameters: zero mismatches allowed (-v 0) and reporting all valid alignments (-a). Mapped reads were sorted and indexed using SAMtools v0.1.1872(53). For small RNA annotation, reads were sequentially mapped to different RNA categories. Structural RNAs (tRNA, rRNA, snRNA, and snoRNA) were annotated based on the Araport11 gene models. miRNA sequences were annotated using 426 mature *Arabidopsis* miRNAs from miRBase(54), including both guide strands (miRNA-5p) and passenger strands (miRNA-3p). Expression levels were quantified as reads per kilobase of transcript per million (RPKM). For genomic loci with multiply mapped reads, RPKM values were calculated as the sum of normalized read counts.

For identification of 21-nt sRNA clusters and selection of control loci, genome-wide small RNA clusters were identified using ShortStack-3.8.5(32) with default parameters. Clusters dominated by 21-nt small RNAs were extracted for further analysis. As a control for phasiRNA abundance ratios, 14 independent randomized sets of non-PHAS 21-nt clusters were randomly selected from the ShortStack-identified clusters were analyzed. One representative result is presented.

### RNA extraction and stem-loop RT-qPCR

Two-week-old *A. thaliana* seedlings grown in liquid MS culture were treated with flg22 or chitin as previously described(55). Total RNA was extracted from *A. thaliana* seedlings using TRIzol reagent and then treated with DNase I to remove genomic DNA contamination. Stem-loop RT-qPCR was conducted following a published protocol(56). 100 ng RNA was used for each reverse transcription reaction. Oligos used for RT-qPCR are listed in Supplementary Table 1.

### Transient protein expression in *N. benthamiana*

Four-week-old *N. benthamiana* plants were used for Agrobacterium-mediated transient expression. Overnight cultures of *Agrobacterium* carrying the constructs of interest were pelleted, resuspended and diluted to OD_600_ = 0.5 in infiltration buffer (10 mM MES, pH 5.6, 10 mM MgCl_2_). Samples for subcellular localization assays and protein expression were harvested 2 days post infiltration.

To examine subcellular localization during pathogen infection, detached leaves were inoculated with the respective pathogens 1 day post infiltration, and the subcellular dynamics of the genes of interest were observed 12 hpi of *P. capsici*, 12 hpi of *P. palmovora*, 60 hpi of C. *higginsianum* and 3 dpi of *P. infestans*. To examine subcellular localization during pathogen infection with flg22 treatment, detached leaves at 2 dpi were submerged in 0.005% Silwet-77 solution with or without 1 μM flg22 peptide for 10 minutes before confocal imaging. Puncta numbers following flg22 treatment were analysed using ImageJ. To assess AGO10 puncta formation following *P. infestans* infection, confocal fields of view were scored for the presence or absence of discrete AGO10 puncta in haustoriated and non-haustoriated regions. For each condition, independent confocal images were analyzed (n = 18 images per condition), with each image treated as a single biological observation to avoid pseudoreplication.

### Co-immunoprecipitation (co-IP) assay

To investigate the interaction between AGO10 and SGS3, FM-AGO10 and SGS3-GFP were transiently co-expressed in *N. benthamiana*. Total proteins were extracted using an extraction buffer (150 mM Tris-HCl, pH7.5, 150 mM NaCl, 1 mM MgCl_2_, 1 mM EDTA, 10% Glycerol, 10 mM DTT, 0.1% NP40, 25 μM MG132, and EDTA-free protease inhibitor cocktail). The protein extracts were immunoprecipitated using pre-equilibrated anti-GFP magnetic beads at 4°C for 1.5 hours. After five washes with the wash buffer (150 mM Tris-HCl, pH7.5, 150 mM NaCl, 1 mM MgCl_2_, 1 mM EDTA, 10% Glycerol, 10 mM DTT, 0.3% NP40), the protein complexes were eluted by heating at 65°C for 15 mins in 2 x SDS loading buffer. AGO10 and SGS3 proteins were detected by immunoblotting using anti-myc and anti-GFP antibody, respectively.

### Fluorescence recovery after photobleaching (FRAP)

FRAP analysis of DsRed-AGO10 condensates was performed on a Leica SP8 laser scanning confocal microscope. The DsRed-AGO10 condensates were photobleached with a 488-nm laser at 100% intensity. Photobleaching was performed at t = 0 s, and fluorescence recovery was monitored every 20 s for a total of 3 minutes post photobleaching. FRAP data analysis was conducted according to methods described by Boeynaems et al(57). FRAP recovery kinetics was quantified by normalizing fluorescence intensity to pre-bleach values. The recovery curve was generated from measurements of 10 independent condensates.

### Phylogenetic analysis of AGO10 proteins

The sequences of AGO10 orthologous proteins were retrieved from the National Center for Biotechnology Information (NCBI) protein database. Multiple sequence alignment was performed using Clustal W in MEGA X software(42). Phylogenetic analysis was conducted using the neighbor-joining method with 1,000 bootstrap replicates. The AGO protein sequence in the AGO1/5/10 clade from the liverwort *Marchantia polymorpha* was designated as the phylogenetic outgroup due to its evolutionary distance from the other plant species analysed. The final phylogenetic tree was visualized and annotated using MEGA X(42).

### Virus-induced gene silencing (VIGS) in *N. benthamiana*

The VIGS vectors *pTRV1* and *pTRV2* were previously described(58). To generate *pTRV2* constructs targeting *GFP*, Nb*AGO10a*, and Nb*AGO10b* genes, around 300-bp fragments from each target gene were PCR-amplified and cloned into *Kpn*I/*Xho*I-digested *pYL156* vectors via homologous recombination. For VIGS assays, Agrobacterium cultures harbouring *pTRV1* were mixed in a 1:1 ratio with cultures containing either *pTRV2-GFP, pTRV2-NbAGO10a*, or *pTRV2-NbAGO10b*, adjusting the final OD600 to 1.0. The bacterial mixtures were infiltrated into young leaves of plants at the 4-leaf stage. Plant phenotypes were monitored for developmental changes over a three-week period post-infiltration and used for pathogen infection.

## Data and materials availability

Plasmids and transgenic plants generated in this study are available from Wenbo Ma upon request under a materials transfer agreement with The Sainsbury Laboratory. The raw data of all sRNA-seq experiments have been deposited in NCBI with project ID PRJNA1215033. All the data is publicly available as of the date of publication. All other data are available in the main text or the supplementary materials.

## Acknowledgments

We thank the technical supports from the cell tissue culture and synthetic biology teams at the Sainsbury Laboratory.

## Funding

Gatsby Charitable Foundation (WM)

UKRI BBSRC Grant BBS/E/J/000PR9797 (WM)

UKRI BBSRC Grant BB/W00691X/1 (WM)

UKRI funded-MSCA Postdoctoral Fellowship (101065015) (MW)

UKRI BBSRC Grant BB/X016382/1 (TOB, ELHY)

UKRI BBSRC Grant BB/T006102/1 (TOB, ELHY)

## Author contributions

Conceptualization: WM

Methodology: MW, XFang, CY, SY, ELHY, BLK, QF, TO, BT, LF, YH

Investigation: MW, XFang, CY, SY, ELHY, BLK, QF, TO, BT, LF

Visualization: MW, XFang, CY, BLK

Project administration: WM

Supervision: WM, XFeng

Writing – original draft: WM, MW

Writing – review & editing: WM, MW, XFeng

## Competing interests

Authors declare that they have no competing interests.

---

**Supplementary figure 1: *A. thaliana ago10* mutants are hypersusceptible to pathogen infection**.

**a**, Disease phenotype of wildtype (Col-0) and *ago* mutants. Four-week-old plants were inoculated with *P. capsici* zoospores by directly applying the zoospore suspension on leaves (indicated with arrowheads). Images were taken at 3 dpi. Scale bars, 2.0 cm. **b**, Schematic representation of the *AGO10* gene structure showing T-DNA insertion sites in two SALK lines. Untranslated regions (UTRs) are colored in grey and exons in blue. **c**, *ago10* mutants were hypersusceptible to the fungal pathogen *Colletotrichum higginsianum*. Leaves of four-week-old plants were inoculated with *C. higginsianum* zoospores and the images showing disease lesions were taken at 5 dpi. Scale bars, 2.0 cm. **d**, Lesion size following *C. higginsianum* infection. Statistical significance was determined using a two-tailed Student’s *t*-test (\*\*\**p* < 0.001). **e**, Catalytic mutants of AGO10 were unable to rescue the hypersusceptibility phenotype of *ago10-1* mutant plants. Four-week-old plants expressing wildtype AGO10 or catalytic mutants (AGO10^D709A^, AGO10^D793A^, and AGO10^H935A^) were inoculated with *P. capsici* zoospores (arrowheads indicate inoculated leaves). Images were taken at 3 dpi. Scale bars, 2.0 cm. **f**, Western blot confirming the expression of AGO10 variants, all tagged with Flag-4Myc at the N-terminus, in transgenic *A. thaliana* lines. Ponceau S staining served as the loading control.

**Supplementary figure 2: AGO10 responds to immune activation at the post-transcription level**.

**a**, *AGO10* transcript level remained unchanged after *P. capsici* infection. Transcript abundance was determined in *A. thaliana* at 5 hpi using qRT-PCR with *ACTIN* as the internal reference. ns represents no significant changes using two-tailed Student’s *t* test. **b**, Single cell transcriptomic data(22) showing transcript levels of *AGO10* in various cell types of *A. thaliana* at 24 and 40 hpi by *C. higginsianum*. **c**, Confirmation of immune activation by flg22 treatment using activation of MPK3/6 as a marker. Western blotting was used to detect phosphorylation of MPK3 and MPK6 using anti-Phospho-p44/42 MAPK (Erk1/2) (Thr202/Tyr204) antibodies. **d**, Chitin treatment induced an increased accumulation of AGO10 proteins. *A. thaliana* plants expressing *p35S::Flag-4Myc-AGO10* were treated with 1 μM chitin and AGO10 proteins were detected using an anti-Myc antibody by western blotting. **e**, AGO1 is not responsive to pathogen infection. AGO1 protein levels were determined using an anti-AGO1 antibody during *P. capsici* infection of *A. thaliana* seedlings. Water served as mock treatment. In **a,c**–**e**, ten-day-old seedlings were used for qRT-PCR or immunoblotting analysis. Ponceau S staining was the loading control in **c**–**e. f**, AGO10 forms cytoplasmic puncta following *P. capsici* infection. DsRed-AGO10 or DsRed were expressed in *N. benthamiana* through Agroinfiltration. Nuclei were visualized by co-expression of H2B-CFP. Scale bars, 25 μm. **g**, Localization of GFP did not change during *P. palmivora* infection. Scale bar,15 μm. **h**, *C. higginsianum* infection induces AGO10 puncta formation in *N. benthamiana*. Images were captured at 3 dpi. Arrowheads indicate the puncta formed in infected cells.

**Supplementary figure 3: AGO10 interacts with SGS3 in siRNA bodies**.

**a**, AGO10-SGS3 interactions were visualized using bimolecular fluorescence complementation (BiFC) in *N. benthamiana* leaves. AGO10 and SGS3 were fused with nYFP and cYFP, respectively. Scale bars, 25 μm. **b**, Structural model of a AGO10-SGS3 interaction interface, which was predicted by AlphaFold Multimer(39) and visualized using ChimeraX(40). Six amino acids of AGO10 (Ile777, Arg780, Lys786, Leu820, Glu821, and Tyr824) were predicted to directly contact with W44 of SGS3, which is within a predicted GW motif. PAE indicates predicted aligned error with low values represent a prediction with high confidence. **c**, Superimposition of the predicted AGO10-SGS3 interaction interfaces with the human hGW182-hAGO1 protein complex (PDB: 4KRE)(28). Models are aligned and visualized using UCSF ChimeraX(40). **d**, AGO10 can no longer form cytoplasmic puncta or co-localize when co-expressed with SGS3^W44A^ in *N. benthamiana*. **e**, AGO10 protein induction during *P. capsici* infection is independent on SGS3. Ten-day-old *A. thaliana* seedlings expressing GFP-AGO10 in either *ago10-1* or *sgs3* mutant background were inoculated with *P. capsici* zoospore. AGO10 protein levels were examined at 5 hpi using anti-GFP antibody by western blotting.

**Supplementary figure 4: AGO10 binds to miRNAs capable of triggering secondary siRNA production after pathogen infection**.

**a**, Reads statistics of sequencing data from AGO10-immunoprecipitated sRNAs. Four-week-old *A. thaliana* transgenic plants expressing *p35S::Flag-4Myc-AGO10* were inoculated with *P. capsici*. AGO10 was immunoprecipitated from *P. capsici*-inoculated or water-treated (mock) samples using an anti-flag beads. Small RNAs co-precipitated with AGO10 were analyzed using sRNA-seq. Three independent replicates were analyzed. **b**, Pearson correlation analysis of 21- and 22-nucleotide miRNA RPKM between biological replicates under mock and *P. capsici* infection conditions. Pearson correlation coefficients (r) are indicated in each panel. **c**, Heatmap showing the relative abundance of 5’ terminal nucleotides in AGO10-bound sRNAs. Data represents percentage of each 5’ nucleotide.

**Supplementary figure 5: AGO10 is required for immune-induced secondary siRNA production**.

**a**, Reads statistics of sequencing data from sRNA analysis of wildtype *A. thaliana* (Col-0) or *ago10-1* mutant plants after flg22 treatment. Ten-day-old seedlings were treated with 1 µm flg22 for 1 or 2 hours before total sRNAs were extracted for sequencing. Three independent replicates were analzyed. **b**, Size distribution of sRNAs in each library showing a predominant peak in 21 nt. **c**, Hierarchical clustering of samples based on Pearson correlation coefficients. Color scale represents Pearson correlation values. **d**, Flg22 treatment induced the accumulation of a secondary siRNA, siR1310 but this induction was abolished in *ago10-1*. The mutant phenotype could be rescued by wildtype but not catalytic mutant of AGO10. The abundance of siR1310 was determined by stemloop-PCR. Different letters indicate statistically significant differences (*p* < 0.05) determined by one-way ANOVA with Tukey’s multiple comparisons test.

**Supplementary figure 6: An N-terminal IDR is required for immune-responsiveness and defense activity of AGO10**.

**a**, Prediction using PONDR(41) suggests an IDR at the N-terminus of AGO10. **b**, Schematic representation of AGO10 protein domain architecture and the construction of AGO10^ΔIDR^ mutant. **c**, Subcellular localization of GFP-AGO10 and GFP-AGO10^ΔIDR^ in *A. thaliana* after *P. capsici* infection. Two-week-old transgenic plants expressing GFP-AGO10 or GFP-AGO10^ΔIDR^ in *ago10-1* background were inoculated with zoospore suspensions or treated with water (mock). Confocal images were taken at 5 hpi. Scale bars, 10 μm. **d**, Western blot confirming the protein accumulation of GFP, GFP-AGO10, or GFP-AGO10^ΔIDR^ in corresponding *A. thaliana* transgenic lines. Proteins were detected using an anti-GFP antibody, with Ponceau S staining as the loading control. **e**, GFP-AGO10^ΔIDR^ was unable to complement the hypersusceptibility phenotype of *ago10-1* mutant. Four-week-old *A. thaliana* plants expressing GFP, GFP-AGO10, or GFP-AGO10^ΔIDR^ were inoculated with *P. capsici* zoospore suspensions. Representative images showing disease symptoms at 3 dpi are presented. Arrowheads indicate inoculated leaves. Scale bars, 2.0 cm.

**Supplementary figure 7: The N-terminal IDR of AGO1 does not respond to pathogen infection**.

**a**, Sequence conservation analysis of the ten *A. thaliana* AGO family members showing the highest level of variation at the N-terminal region. **b**, Prediction using PONDR(41) suggests an IDR at the N-terminus of *A. thaliana* AGO1. **c**, Sequence alignment of *A. thaliana* AGO10 and AGO1 using ClustalW in MEGA X(42). The N-terminal regions (with yellow-colored underline) indicate swapped sequences used to generate the chimeric constructs AGO1^AGO10-IDR^ and AGO10^AGO1-IDR^. **d**, A schematic showing the construction of chimeric AGO1 and AGO10 proteins with their N-terminal IDR regions swapped. **e**, Images of four-week-old transgenic *A. thaliana* expressing AGO1 or the AGO1^AGO10-IDR^ chimera in *ago10-1* mutant background. **f**, Western blotting confirming the protein accumulation of AGO1 or AGO1^AGO10-IDR^ in the transgenic lines. Protein accumulation was detected by immunoblotting using an anti-GFP antibody, with Ponceau S staining as the loading control. **g**, AGO1 or the AGO1^AGO10-IDR^ did not rescue the *ago10-1* mutant phenotype in plant defense. Four-week-old *A. thaliana* plants were inoculated with *P. capsici* zoospores. Pathogen biomass was determined at 3 dpi. Different letters indicate significant differences (*p* < 0.05, one-way ANOVA with Tukey’s multiple comparisons).

**Supplementary figure 8: A conserved function of AGO10a subclade members in plant immunity**.

**a**, Phylogenetic analysis of AGO10 homologs showing two subclades – AGO10a and AGO10b. AGO10 homologs from representative plant species, including *A. thaliana* (At), *Glycine max* (Glyma), *Marchantia polymorpha* (Mapoly), *N. benthamiana* (Nb), *Nicotiana tabacum* (Nt), *Oryza sativa* (Os), and *Solanum lycopersicum* (Solyc), were analyzed. The liverwort Marchantia encodes one protein in the AGO1/5/10 clade (Mapoly0001s0149 or MapolyAGO in the tree), which was used as the outgroup. The phylogenetic tree was constructed using the neighbour-joining method in MEGA X(42) with 1,000 bootstrap replicates. **b**, Distribution of proline residues in the N-terminal region shows distinct patterns in AGO10a and AGO10b subclades, which are indicated by blue and red, respectively. The yellow line represents MapolyAGO, and the dashed line represents OsAGO10. The proline ratio was calculated using a sliding window of 10 amino acids with a 5-amino-acid step size. **c**, Multiple sequence alignment of AGO10 proteins from different plant species using ClustalW(43) shows a high-level variation in the N-terminal region. **d**, An enlarged view of the N-terminal region highlighting differences in the predicted proline-rich domain (PRD) between AGO10a and AGO10b proteins. **e**, *NbAGO10a* and *NbAGO10b* were efficiently silenced in *N. benthamiana* using VIGS. Transcript abundances were analysed by RT-qPCR with *NbEF1α* as the internal reference. The VIGS vector carrying *GFP* was used as the negative control. Data represent means from three independent experiments ± SEM. Statistical significance was determined using two-tailed Student’s t-test (\*\**p* < 0.01; ns, not significant). **f**, *NbAGO10a*- and *NbAGO10b*-silenced *N. benthamiana* plants exhibited normal growth. Representative images of plants subjected to VIGS targeting *NbAGO10a* or *NbAGO10b*, respectively. Plants infected with the VIGS vector carrying GFP were used as the control. Scale bar, 5.0 cm.

## Notes

### Competing Interest Statement

The authors have declared no competing interest.

## References

1. X. Chen, O. Rechavi, Plant and animal small RNA communications between cells and organisms. Nat. Rev. Mol. Cell Biol. 23, 185–203 (2022).

2. J. Zhan, B. C. Meyers, Plant small RNAs: Their biogenesis, regulatory roles, and functions. Annu. Rev. Plant Biol. 74, 21–51 (2023).

3. Y. Qiao et al., Small RNAs in plant immunity and virulence of filamentous pathogens. Annu. Rev. Phytopathol. 59, 265–288 (2021).

4. Y. Hou, W. Ma, Natural host-induced gene silencing offers new opportunities to engineer disease resistance. Trends Microbiol. 28, 109–117 (2020).

5. H. Zand Karimi, R. W. Innes, Molecular mechanisms underlying host-induced gene silencing. The Plant Cell 34, 3183–3199 (2022).

6. Y. Yu, H. Wang, C. You, X. Chen, Plant microRNA maturation and function. Nat. Rev. Mol. Cell Biol. 27, 55–70 (2026).

7. M. J. Axtell, C. Jan, R. Rajagopalan, D. P. Bartel, A two-hit trigger for siRNA biogenesis in plants. Cell 127, 565–577 (2006).

8. H. Vaucheret, Plant ARGONAUTES. Trends Plant Sci. 13, 350–358 (2008).

9. Q. Fei, R. Xia, B. C. Meyers, Phased, secondary, small interfering RNAs in posttranscriptional regulatory networks. Plant Cell 25, 2400–2415 (2013).

10. U. Ellendorff, E. F. Fradin, R. de Jonge, B. P. H. J. Thomma, RNA silencing is required for Arabidopsis defence against Verticillium wilt disease. J. Exp. Bot. 60, 591–602 (2008).

11. Y. Hou et al., A Phytophthora effector suppresses trans-kingdom RNAi to promote disease susceptibility. Cell Host Microbe 25, 153–165 (2019).

12. Q. Cai et al., Plants send small RNAs in extracellular vesicles to fungal pathogen to silence virulence genes. Science 360, 1126–1129 (2018).

13. Y. Qiao et al., Oomycete pathogens encode RNA silencing suppressors. Nat. Genet. 45, 330–333 (2013).

14. C. Yin et al., A novel fungal effector from Puccinia graminis suppressing RNA silencing and plant defense responses. New Phytol. 222, 1561–1572 (2019).

15. H. Zhu et al., Arabidopsis Argonaute10 specifically sequesters mir166/165 to regulate shoot apical meristem development. Cell 145, 242–256 (2011).

16. Y. Xiao, S. Maeda, T. Otomo, I. J. MacRae, Structural basis for RNA slicing by a plant Argonaute. Nat. Struct. Mol. Biol. 30, 778–784 (2023).

17. Y. Yu et al., ARGONAUTE10 promotes the degradation of miR165/6 through the SDN1 and SDN2 exonucleases in Arabidopsis. PLoS Biol. 15, e2001272 (2017).

18. Y. Zhou et al., Spatiotemporal sequestration of mir165/166 by Arabidopsis Argonaute10 promotes shoot apical meristem maintenance. Cell Rep. 10, 1819–1827 (2015).

19. N. El Arbi et al., ARGONAUTE10 controls cell fate specification and formative cell divisions in the Arabidopsis root. EMBO J. 43, 1822–1842 (2024).

20. S. Mirlohi, G. Schott, A. Imboden, O. Voinnet, An AGO10:miR165/6 module regulates meristem activity and xylem development in the Arabidopsis root. EMBO J. 43, 1843–1869 (2024).

21. L. Feng et al., A conserved small RNA-generating gene cluster undergoes sequence diversification and contributes to plant immunity. bioRxiv 10.1101/2025.07.20.665670, 2025.2007.2020.665670 (2025).

22. B. Tang, L. Feng, M. T. Hulin, P. Ding, W. Ma, Cell-type-specific responses to fungal infection in plants revealed by single-cell transcriptomics. Cell Host Microbe 31, 1732–1747 (2023).

23. C. Duggan et al., Dynamic localization of a helper NLR at the plant–pathogen interface underpins pathogen recognition. Proc. Natl. Acad. Sci. U. S. A. 118, e2104997118 (2021).

24. N. Kumakura et al., SGS3 and RDR6 interact and colocalize in cytoplasmic SGS3/RDR6-bodies. FEBS Lett. 583, 1261–1266 (2009).

25. H. Tan et al., Phase separation of SGS3 drives siRNA body formation and promotes endogenous gene silencing. Cell Rep. 42, 111985 (2023).

26. R. Evans et al., Protein complex prediction with AlphaFold-Multimer. bioRxiv 10.1101/2021.10.04.463034 (2022).

27. M. El-Shami et al., Reiterated WG/GW motifs form functionally and evolutionarily conserved ARGONAUTE-binding platforms in RNAi-related components. Genes Dev. 21, 2539–2544 (2007).

28. Christopher R. Faehnle, E. Elkayam, Astrid D. Haase, Gregory J. Hannon, L. Joshua-Tor, The making of a slicer: Activation of human Argonaute-1. Cell Rep. 3, 1901–1909 (2013).

29. X. Fang, Y. Qi, RNAi in plants: An Argonaute-centered view. The Plant Cell 28, 272–285 (2016).

30. P. A. Manavella, D. Koenig, D. Weigel, Plant secondary siRNA production determined by microRNA-duplex structure. Proc. Natl. Acad. Sci. U. S. A. 109, 2461–2466 (2012).

31. C. Chen et al., sRNAanno—a database repository of uniformly annotated small RNAs in plants. Horticulture Research 8, 45 (2021).

32. N. R. Johnson, J. M. Yeoh, C. Coruh, M. J. Axtell, Improved Placement of Multi-mapping Small RNAs. G3 Genes|Genomes|Genetics 6, 2103–2111 (2016).

33. A. S. Holehouse, B. B. Kragelund, The molecular basis for cellular function of intrinsically disordered protein regions. Nat. Rev. Mol. Cell Biol. 25, 187–211 (2024).

34. A. Blagojevic et al., Heat stress promotes Arabidopsis AGO1 phase separation and association with stress granule components. iScience 27, 109151 (2024).

35. H. Narita, T. Shima, R. Iizuka, S. Uemura, N-terminal region of Drosophila melanogaster Argonaute2 forms amyloid-like aggregates. BMC Biol. 21, 78 (2023).

36. S. Bélanger, J. Zhan, B. C. Meyers, Phylogenetic analyses of seven protein families refine the evolution of small RNA pathways in green plants. Plant Physiol. 192, 1183–1203 (2023).

37. M. Yoshikawa et al., Cooperative recruitment of RDR6 by SGS3 and SDE5 during small interfering RNA amplification in Arabidopsis. Proc. Natl. Acad. Sci. U. S. A. 118, e2102885118 (2021).

38. Y. Xiao, I. J. MacRae, The molecular mechanism of microRNA duplex selectivity of Arabidopsis ARGONAUTE10. Nucleic Acids Res. 50, 10041–10052 (2022).

39. J. Jumper et al., Highly accurate protein structure prediction with AlphaFold. Nature 596, 583–589 (2021).

40. E. C. Meng et al., UCSF ChimeraX: Tools for structure building and analysis. Protein Sci. 32, e4792 (2023).

41. P. Romero et al., Sequence complexity of disordered protein. Proteins: Structure, Function, and Bioinformatics 42, 38–48 (2001).

42. S. Kumar, G. Stecher, M. Li, C. Knyaz, K. Tamura, MEGA X: Molecular evolutionary genetics analysis across computing platforms. Mol. Biol. Evol. 35, 1547–1549 (2018).

43. J. D. Thompson, D. G. Higgins, T. J. Gibson, CLUSTAL W: improving the sensitivity of progressive multiple sequence alignment through sequence weighting, position-specific gap penalties and weight matrix choice. Nucleic Acids Res. 22, 4673–4680 (1994).

44. Y. Wang, K. Bouwmeester, J. E. van de mortel, W. Shan, F. Govers, A novel Arabidopsis–oomycete pathosystem: differential interactions with Phytophthora capsici reveal a role for camalexin, indole glucosinolates and salicylic acid in defence. Plant Cell Environ. 36, 1192–1203 (2013).

45. R. O’Connell et al., A novel Arabidopsis-Colletotrichum pathosystem for the molecular dissection of plant-fungal interactions. MPMI. 17, 272–282 (2004).

46. P. Carella, A. Gogleva, M. Tomaselli, C. Alfs, S. Schornack, Phytophthora palmivora establishes tissue-specific intracellular infection structures in the earliest divergent land plant lineage. Proc. Natl. Acad. Sci. U. S. A. 115, 3846–3855 (2018).

47. J. Long et al., Nurse cell-derived small RNAs define paternal epigenetic inheritance in Arabidopsis. Science 373, eabh0556 (2021).

48. L. Feng et al., An online database for exploring over 2,000 Arabidopsis small RNA libraries. Plant Physiol. 182, 685–691 (2019).

49. J. Tsuji, Z. Weng, DNApi: A De Novo adapter prediction algorithm for small RNA sequencing data. PLoS One 11, e0164228 (2016).

50. M. Martin, Cutadapt removes adapter sequences from high-throughput sequencing reads. EMBnet J. 17, 3 (2011).

51. C. Y. Cheng et al., Araport11: A complete reannotation of the Arabidopsis thaliana reference genome. Plant J. 89, 789–804 (2017).

52. B. Langmead, C. Trapnell, M. Pop, S. L. Salzberg, Ultrafast and memory-efficient alignment of short DNA sequences to the human genome. Genome Biol. 10, R25 (2009).

53. H. Li et al., The Sequence Alignment/Map format and SAMtools. Bioinformatics 25, 2078–2079 (2009).

54. A. Kozomara, M. Birgaoanu, S. Griffiths-Jones, miRBase: from microRNA sequences to function. Nucleic Acids Res. 47, 155–162 (2018).

55. M. Bjornson, P. Pimprikar, T. Nürnberger, C. Zipfel, The transcriptional landscape of Arabidopsis thaliana pattern-triggered immunity. Nat. Plants 7, 579–586 (2021).

56. E. Varkonyi-Gasic, R. Wu, M. Wood, E. F. Walton, R. P. Hellens, Protocol: A highly sensitive RT-PCR method for detection and quantification of microRNAs. Plant Methods 3, 12 (2007).

57. S. Boeynaems, M. De Decker, P. Tompa, L. Van Den Bosch, Arginine-rich peptides can actively mediate liquid-liquid phase separation. Bio Protoc. 7, e2525 (2017).

58. A. C. Velásquez, S. Chakravarthy, G. B. Martin, Virus-induced Gene Silencing (VIGS) in Nicotiana benthamiana and Tomato. Vis. Exp. 28, e1292 (2009).

