## Supplementary figures for "A specialized ARGONAUTE enables trans-species RNA interference in plant immunity"

#### Supplementary figure 1

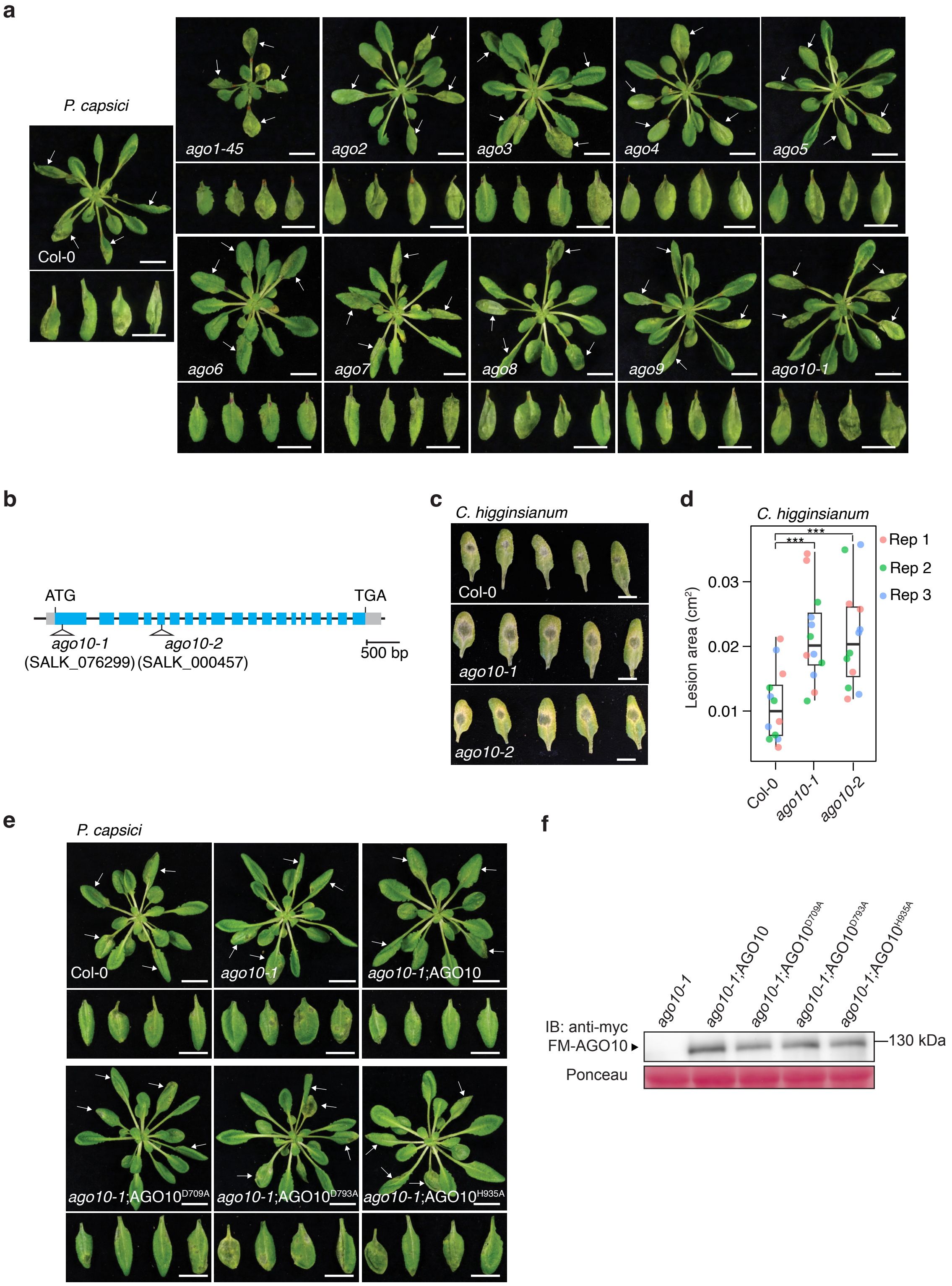


**Supplementary figure1: *A. thaliana ago10* mutants are hypersusceptible to pathogen infection.**
**a,** Disease phenotype of wildtype (Col-0) and *ago* mutants. Four-week-old plants were inoculated with *P. capsici* zoospores by directly applying the zoospore suspension on leaves (indicated with arrowheads). Images were taken at 3 dpi. Scale bars, 2.0 cm. **b,** Schematic representation of the *AGO10* gene structure showing T-DNA insertion sites in two SALK lines. Untranslated regions (UTRs) are colored in grey and exons in blue. **c,** *ago10* mutants were hypersusceptible to the fungal pathogen *Colletotrichum higginsianum*. Leaves of four-week-old plants were inoculated with *C. higginsianum* zoospores and the images showing disease lesions were taken at 5 dpi. Scale bars, 2.0 cm. **d,** Lesion size following *C. higginsianum* infection. Statistical significance was determined using a two-tailed Student's *t*-test (***$p < 0.001$). **e,** Catalytic mutants of AGO10 were unable to rescue the hypersusceptibility phenotype of *ago10-1* mutant plants. Four-week-old plants expressing wildtype AGO10 or catalytic mutants ($AGO10^{D709A}$, $AGO10^{D793A}$, and $AGO10^{H935A}$) were inoculated with *P. capsici* zoospores (arrowheads indicate inoculated leaves). Images were taken at 3 dpi. Scale bars, 2.0 cm. **f,** Western blot confirming the expression of AGO10 variants, all tagged with Flag-4Myc at the N-terminus, in transgenic *A. thaliana* lines. Ponceau S staining served as the loading control.

**Supplementary figure 2**

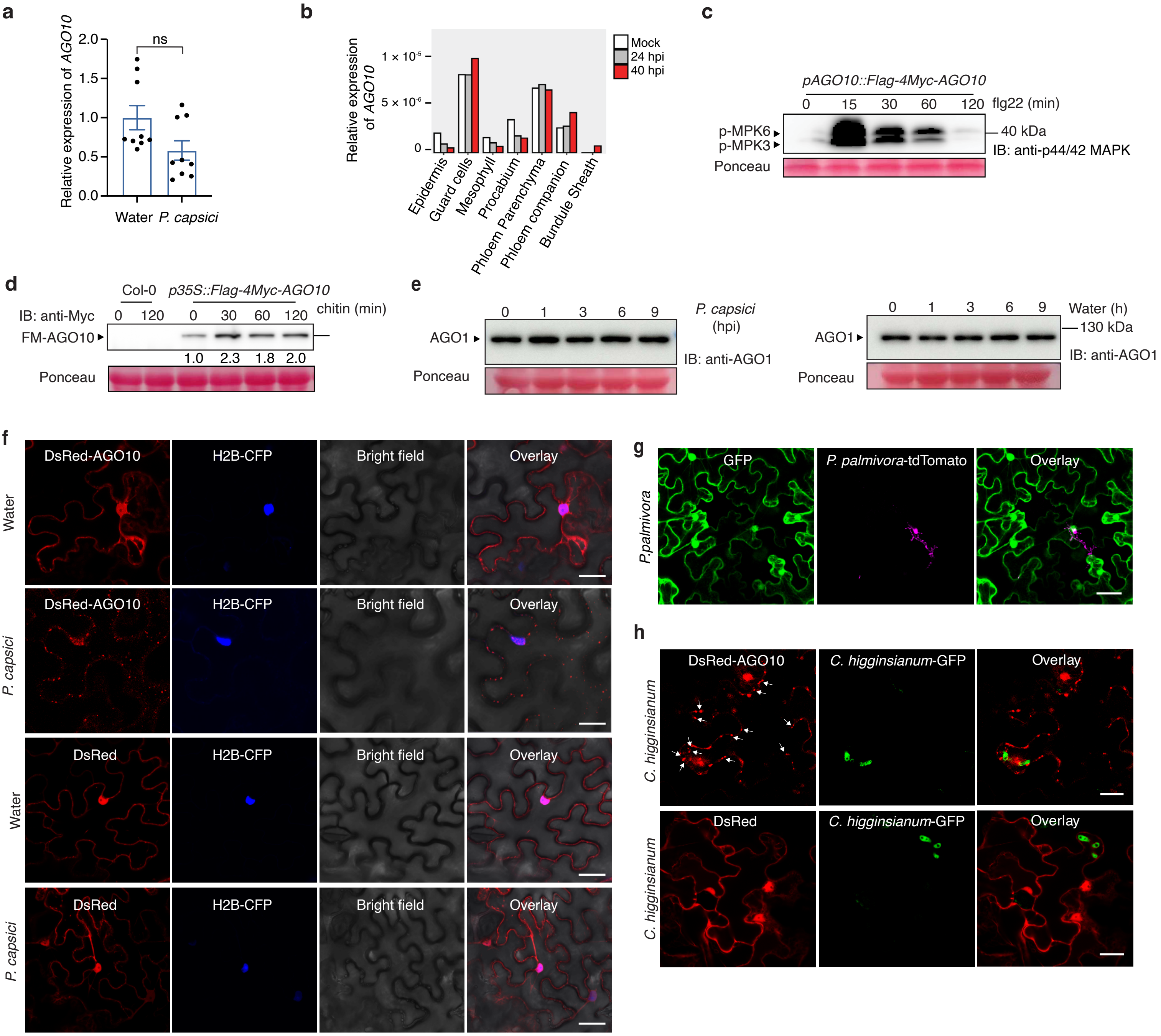


**Supplementary figure2: AGO10 responds to immune activation at the post-transcription level.**
**a,** *AGO10* transcript level remained unchanged after *P. capsici* infection. Transcript abundance was determined in *A. thaliana* at 5 hpi using qRT-PCR with *ACTIN* as the internal reference. ns represents no significant changes using two-tailed Student's *t* test. **b,** Single cell transcriptomic data(22) showing transcript levels of *AGO10* in various cell types of *A. thaliana* at 24 and 40 hpi by *C. higginsianum*. **c**, Confirmation of immune activation by flg22 treatment using activation of MPK3/6 as a marker. Western blotting was used to detect phosphorylation of MPK3 and MPK6 using anti-Phospho-p44/42 MAPK (Erk1/2) (Thr202/Tyr204) antibodies. **d**, Chitin treatment induced an increased accumulation of AGO10 proteins. *A. thaliana* plants expressing *p35S::Flag-4Myc-AGO10* were treated with 1 µM chitin and AGO10 proteins were detected using an anti-Myc antibody by western blotting. **e**, AGO1 is not responsive to pathogen infection. AGO1 protein levels were determined using an anti-AGO1 antibody during *P. capsici* infection of *A. thaliana* seedlings. Water served as mock treatment. In **a,c–e**, ten-day-old seedlings were used for qRT-PCR or immunoblotting analysis. Ponceau S staining was the loading control in **c–e**. **f,** AGO10 forms cytoplasmic puncta following *P. capsici* infection. DsRed-AGO10 or DsRed were expressed in *N. benthamiana* through Agroinfiltration. Nuclei were visualized by co-expression of H2B-CFP. Scale bars, 25 µm. **g,** Localization of GFP did not change during *P. palmivora* infection. Scale bar,15 µm. **h,** *C. higginsianum* infection induces AGO10 puncta formation in *N. benthamiana*. Images were captured at 3 dpi. Arrowheads indicate the puncta formed in infected cells.

**Supplementary figure 3**

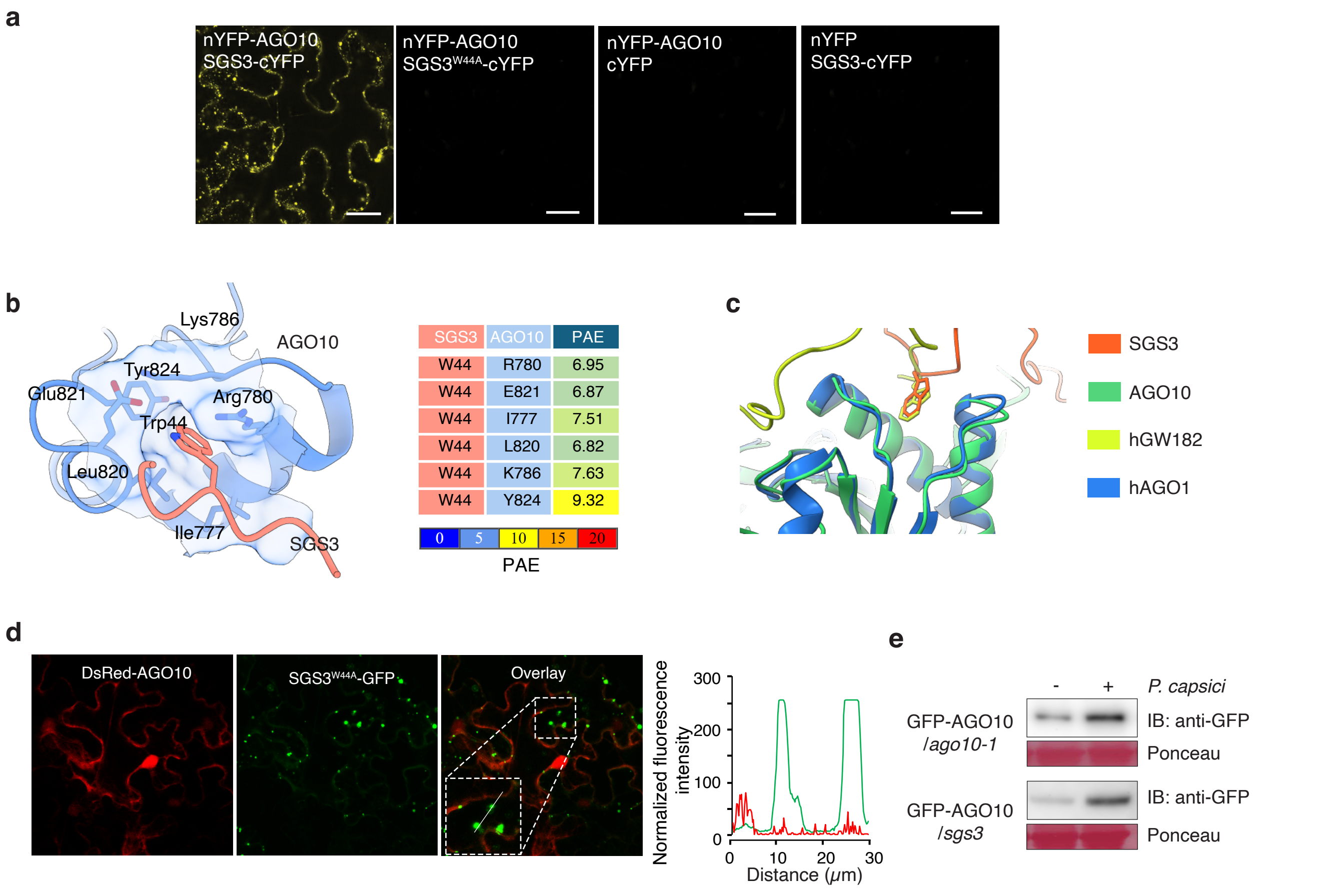


**Supplementary figure3: AGO10 interacts with SGS3 in siRNA bodies.**

**a,** AGO10-SGS3 interactions were visualized using bimolecular fluorescence complementation (BiFC) in *N. benthamiana* leaves. AGO10 and SGS3 were fused with nYFP and cYFP, respectively. Scale bars, 25 µm. **b,** Structural model of a AGO10-SGS3 interaction interface, which was predicted by AlphaFold Multimer(39) and visualized using ChimeraX(40). Six amino acids of AGO10 (Ile777, Arg780, Lys786, Leu820, Glu821, and Tyr824) were predicted to directly contact with W44 of SGS3, which is within a predicted GW motif. PAE indicates predicted aligned error with low values represent a prediction with high confidence. **c,** Superimposition of the predicted AGO10-SGS3 interaction interfaces with the human hGW182-hAGO1 protein complex (PDB: 4KRE)(28). Models are aligned and visualized using UCSF ChimeraX(40). **d,** AGO10 can no longer form cytoplasmic puncta or co-localize when co-expressed with SGS3$^{W44A}$ in *N. benthamiana*. **e,** AGO10 protein induction during *P. capsici* infection is independent on SGS3. Ten-day-old *A. thaliana* seedlings expressing GFP-AGO10 in either *ago10-1* or *sgs3* mutant background were inoculated with *P. capsici* zoospore. AGO10 protein levels were examined at 5 hpi using anti-GFP antibody by western blotting.

**Supplementary figure 4**

**a**

| Libraries | Clean reads | Mapped reads |
|---|---|---|
| AGO10IP_mock_rep1 | 4,829,932 | 1,455,523 (30.14%) |
| AGO10IP_mock_rep2 | 8,980,697 | 4,444,318 (49.49%) |
| AGO10IP_mock_rep3 | 14,271,937 | 7,512,734 (52.64%) |
| AGO10IP_*P.capsici*_rep1 | 15,026,491 | 8,939,380 (59.49%) |
| AGO10IP_*P.capsici*_rep2 | 21,430,531 | 15,409,884 (71.91%) |
| AGO10IP_*P.capsici*_rep3 | 22,527,722 | 15,942,683 (70.77%) |

**c**

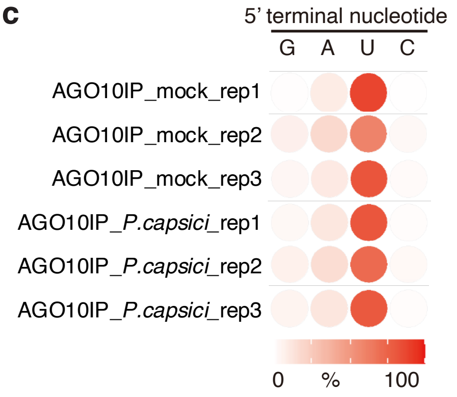


**b**

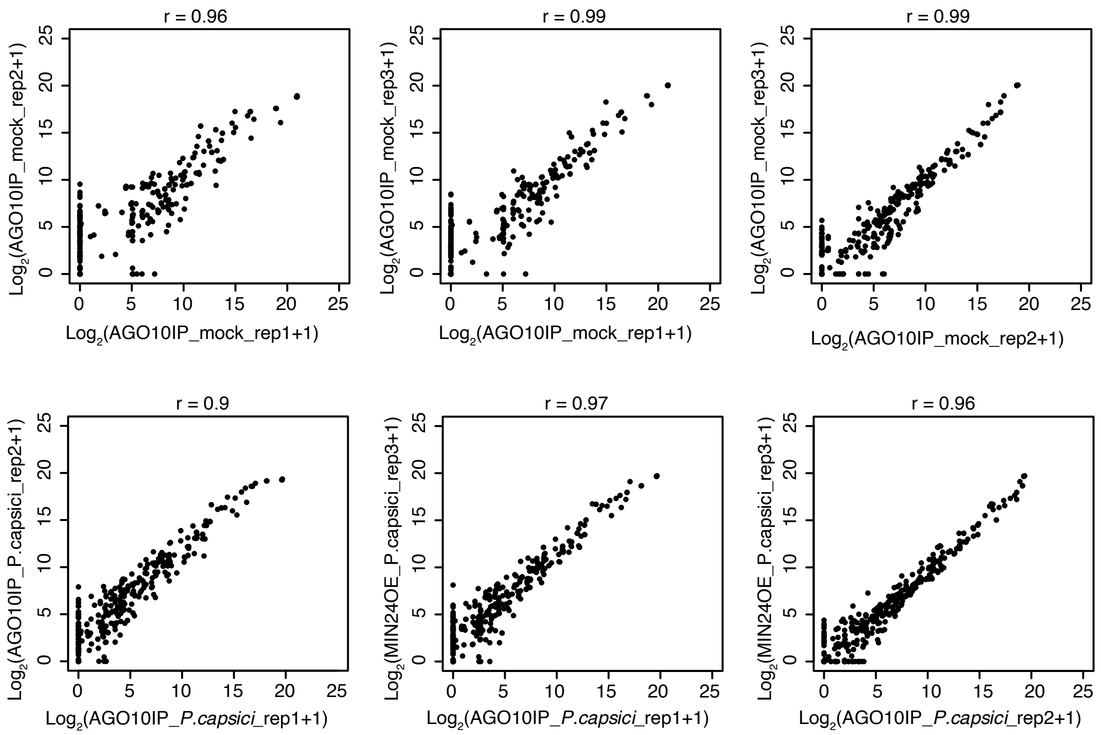


**Supplementary figure 4: AGO10 binds to miRNAs capable of triggering secondary siRNA production after pathogen infection.**

**a,** Reads statistics of sequencing data from AGO10-immunoprecipitated sRNAs. Four-week-old *A. thaliana* transgenic plants expressing *p35S::Flag-4Myc-AGO10* were inoculated with *P. capsici*. AGO10 was immunoprecipitated from *P. capsici*-inoculated or water-treated (mock) samples using an anti-flag beads. Small RNAs co-precipitated with AGO10 were analyzed using sRNA-seq. Three independent replicates were analyzed. **b,** Pearson correlation analysis of 21- and 22-nucleotide miRNA RPKM between biological replicates under mock and *P. capsici* infection conditions. Pearson correlation coefficients (r) are indicated in each panel. **c,** Heatmap showing the relative abundance of 5' terminal nucleotides in AGO10-bound sRNAs. Data represents percentage of each 5' nucleotide.

#### Supplementary figure 5

**a**

| Library Name | Total reads | Mapped Reads | Mapping_rate |
|---|---|---|---|
| Col0_0hrep1 | 3,778,103 | 3,267,827 | 86.49% |
| Col0_1hrep1 | 2,819,570 | 2,423,494 | 85.95% |
| Col0_2hrep1 | 2,134,831 | 1,836,041 | 86.00% |
| *ago10*_0hrep1 | 4,401,244 | 3,853,671 | 87.56% |
| *ago10*_1hrep1 | 7,252,234 | 6,270,010 | 86.46% |
| *ago10*_2hrep1 | 2,117,779 | 1,826,918 | 86.27% |
| Col0_0hrep2 | 4,293,665 | 3,706,813 | 86.33% |
| Col0_1hrep2 | 3,173,609 | 2,731,380 | 86.07% |
| Col0_2hrep2 | 6,246,489 | 5,509,120 | 88.20% |
| *ago10*_0hrep2 | 4,614,853 | 4,005,487 | 86.80% |
| *ago10*_1hrep2 | 3,956,803 | 3,408,965 | 86.15% |
| *ago10*_2hrep2 | 3,616,297 | 3,113,250 | 86.09% |
| Col0_0hrep3 | 4,502,577 | 3,859,875 | 85.73% |
| Col0_1hrep3 | 5,532,670 | 4,757,306 | 85.99% |
| Col0_2hrep3 | 8,872,749 | 7,527,114 | 84.83% |
| *ago10*_0hrep3 | 4,375,624 | 3,762,216 | 85.98% |
| *ago10*_1hrep3 | 4,117,952 | 3,546,064 | 86.11% |
| *ago10*_2hrep3 | 4,084,018 | 3,478,069 | 85.16% |

**b**

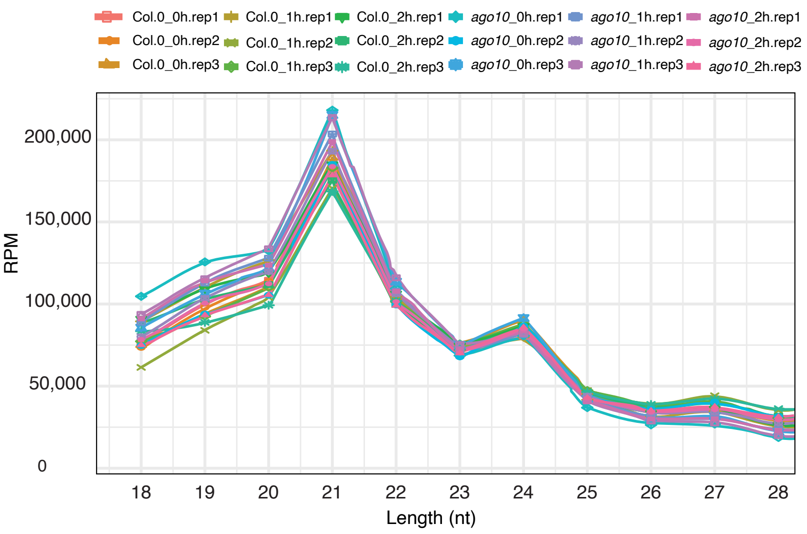


**c** **d**

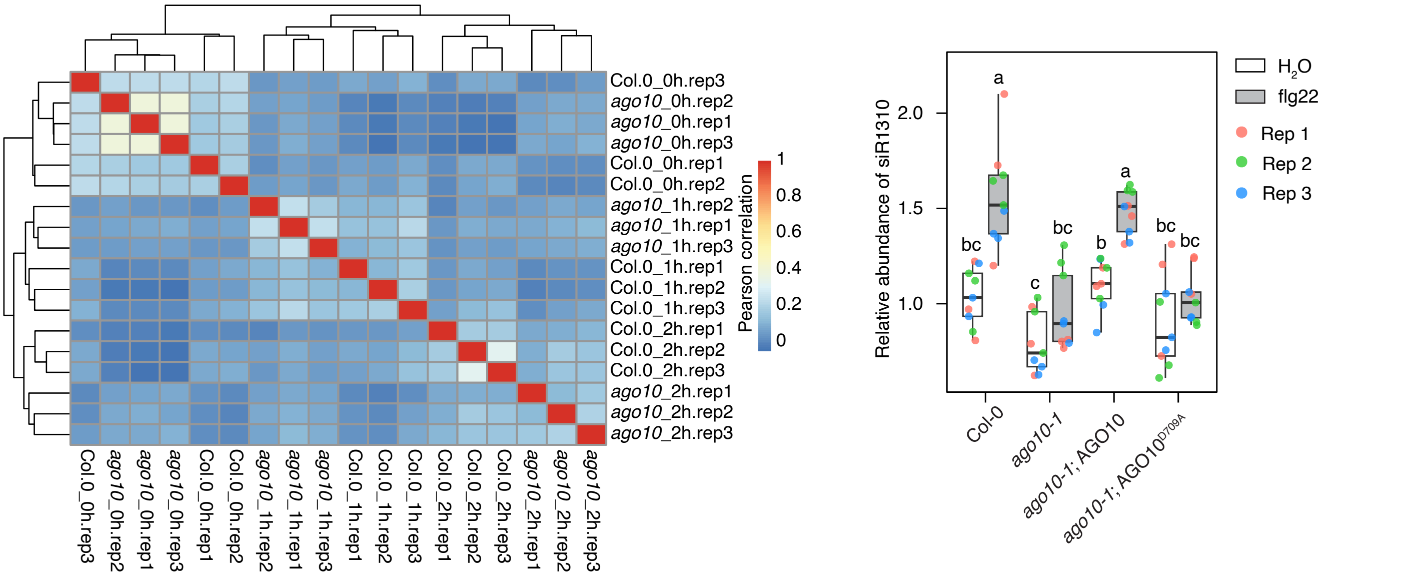


**Supplementary figure 5: AGO10 is required for immune-induced secondary siRNA production.**
**a**, Reads statistics of sequencing data from sRNA analysis of wildtype *A. thaliana* (Col-0) or *ago10-1* mutant plants after flg22 treatment. Ten-day-old seedlings were treated with 1 $\mu$m flg22 for 1 or 2 hours before total sRNAs were extracted for sequencing. Three independent replicates were analzyed. **b**, Size distribution of sRNAs in each library showing a predominant peak in 21 nt. **c**, Hierarchical clustering of samples based on Pearson correlation coefficients. Color scale represents Pearson correlation values. **d**, Flg22 treatment induced the accumulation of a secondary siRNA, siR1310 but this induction was abolished in *ago10-1*. The mutant phenotype could be rescued by wildtype but not catalytic mutant of AGO10. The abundance of siR1310 was determined by stemloop-PCR. Different letters indicate statistically significant differences ($p < 0.05$) determined by one-way ANOVA with Tukey's multiple comparisons test.

#### Supplementary figure 6

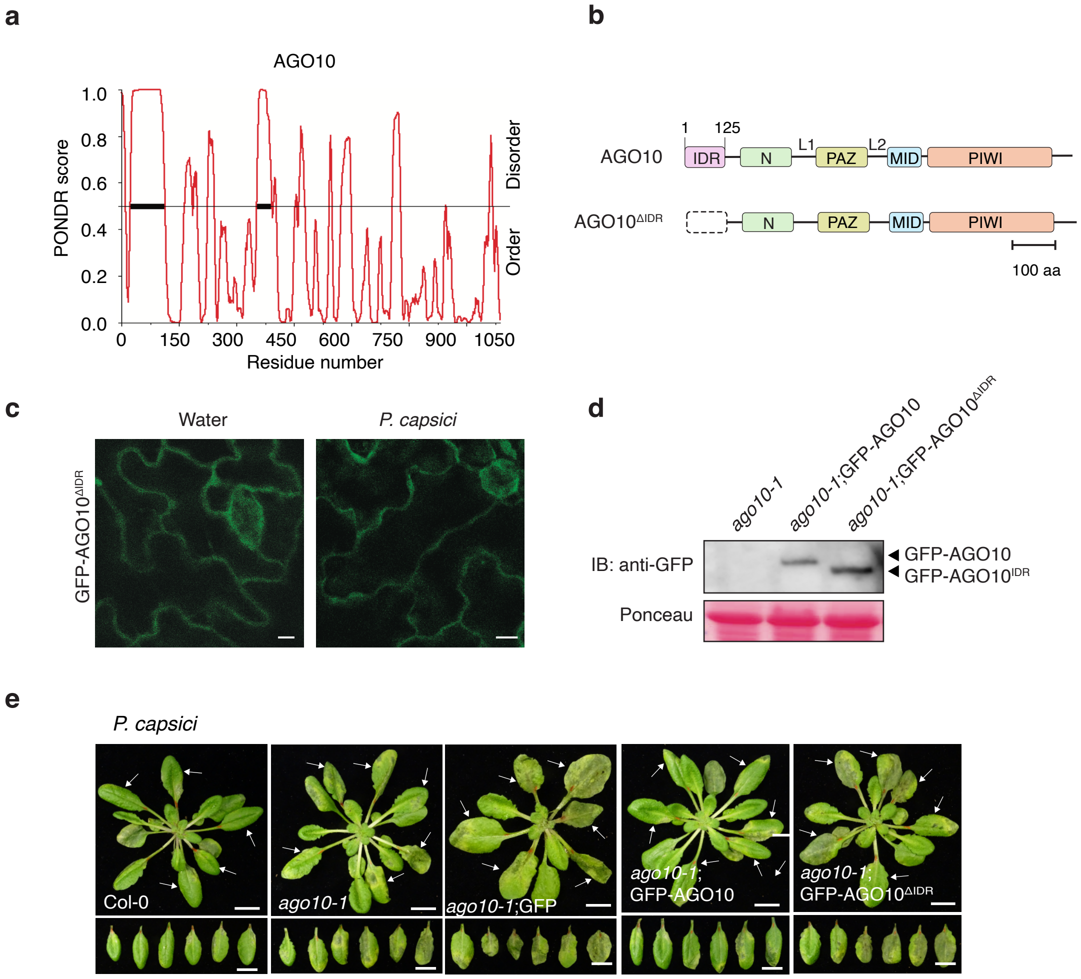


**Supplementary figure 6: An N-terminal IDR is required for immune-responsiveness and defense activity of AGO10.**

**a,** Prediction using PONDR(41) suggests an IDR at the N-terminus of AGO10. **b,** Schematic representation of AGO10 protein domain architecture and the construction of $AGO10^{\Delta IDR}$ mutant. **c,** Subcellular localization of GFP-AGO10 and GFP-$AGO10^{\Delta IDR}$ in *A. thaliana* after *P. capsici* infection. Two-week-old transgenic plants expressing GFP-AGO10 or GFP-$AGO10^{\Delta IDR}$ in *ago10-1* background were inoculated with zoospore suspensions or treated with water (mock). Confocal images were taken at 5 hpi. Scale bars, 10 µm. **d,** Western blot confirming the protein accumulation of GFP, GFP-AGO10, or GFP-$AGO10^{\Delta IDR}$ in corresponding *A. thaliana* transgenic lines. Proteins were detected using an anti-GFP antibody, with Ponceau S staining as the loading control. **e,** GFP-$AGO10^{\Delta IDR}$ was unable to complement the hypersusceptibility phenotype of *ago10-1* mutant. Four-week-old *A. thaliana* plants expressing GFP, GFP-AGO10, or GFP-$AGO10^{\Delta IDR}$ were inoculated with *P. capsici* zoospore suspensions. Representative images showing disease symptoms at 3 dpi are presented. Arrowheads indicate inoculated leaves. Scale bars, 2.0 cm.

### Supplementary figure 7

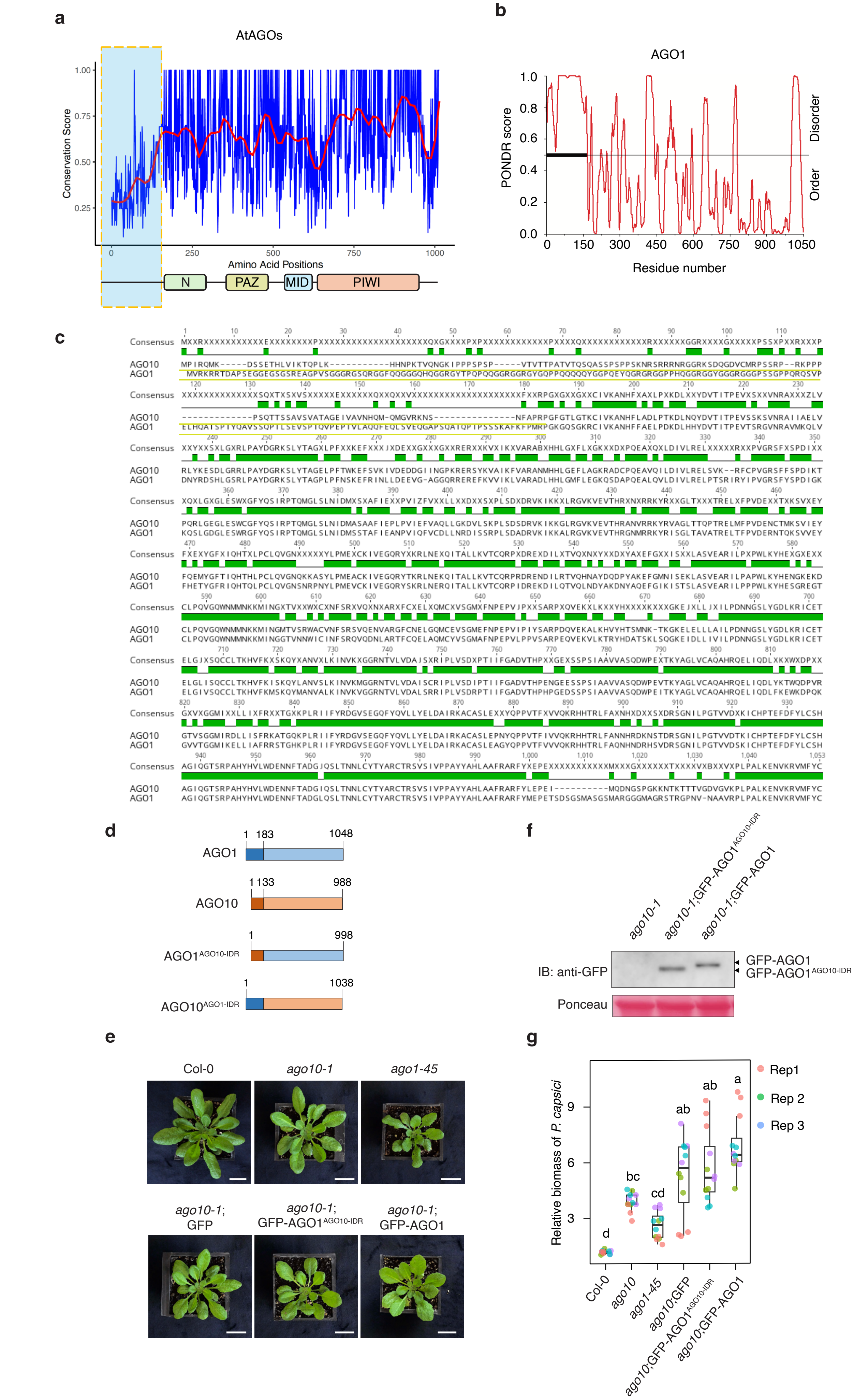

**Supplementary figure 7: The N-terminal IDR of AGO1 does not respond to pathogen infection.**

**a,** Sequence conservation analysis of the ten *A. thaliana* AGO family members showing the highest level of variation at the N-terminal region. **b,** Prediction using PONDR(41) suggests an IDR at the N-terminus of *A. thaliana* AGO1. **c,** Sequence alignment of *A. thaliana* AGO10 and AGO1 using ClustalW in MEGA X(42). The N-terminal regions (with yellow-colored underline) indicate swapped sequences used to generate the chimeric constructs $AGO1^{AGO10\text{-}IDR}$ and $AGO10^{AGO1\text{-}IDR}$. **d**, A schematic showing the construction of chimeric AGO1 and AGO10 proteins with their N-terminal IDR regions swapped. **e**, Images of four-week-old transgenic *A. thaliana* expressing AGO1 or the $AGO1^{AGO10\text{-}IDR}$ chimera in *ago10-1* mutant background. **f**, Western blotting confirming the protein accumulation of AGO1 or $AGO1^{AGO10\text{-}IDR}$ in the transgenic lines. Protein accumulation was detected by immunoblotting using an anti-GFP antibody, with Ponceau S staining as the loading control. **g**, AGO1 or the $AGO1^{AGO10\text{-}IDR}$ did not rescue the *ago10-1* mutant phenotype in plant defense. Four-week-old *A. thaliana* plants were inoculated with *P. capsici* zoospores. Pathogen biomass was determined at 3 dpi. Different letters indicate significant differences ($p < 0.05$, one-way ANOVA with Tukey's multiple comparisons).

#### Supplementary figure 8

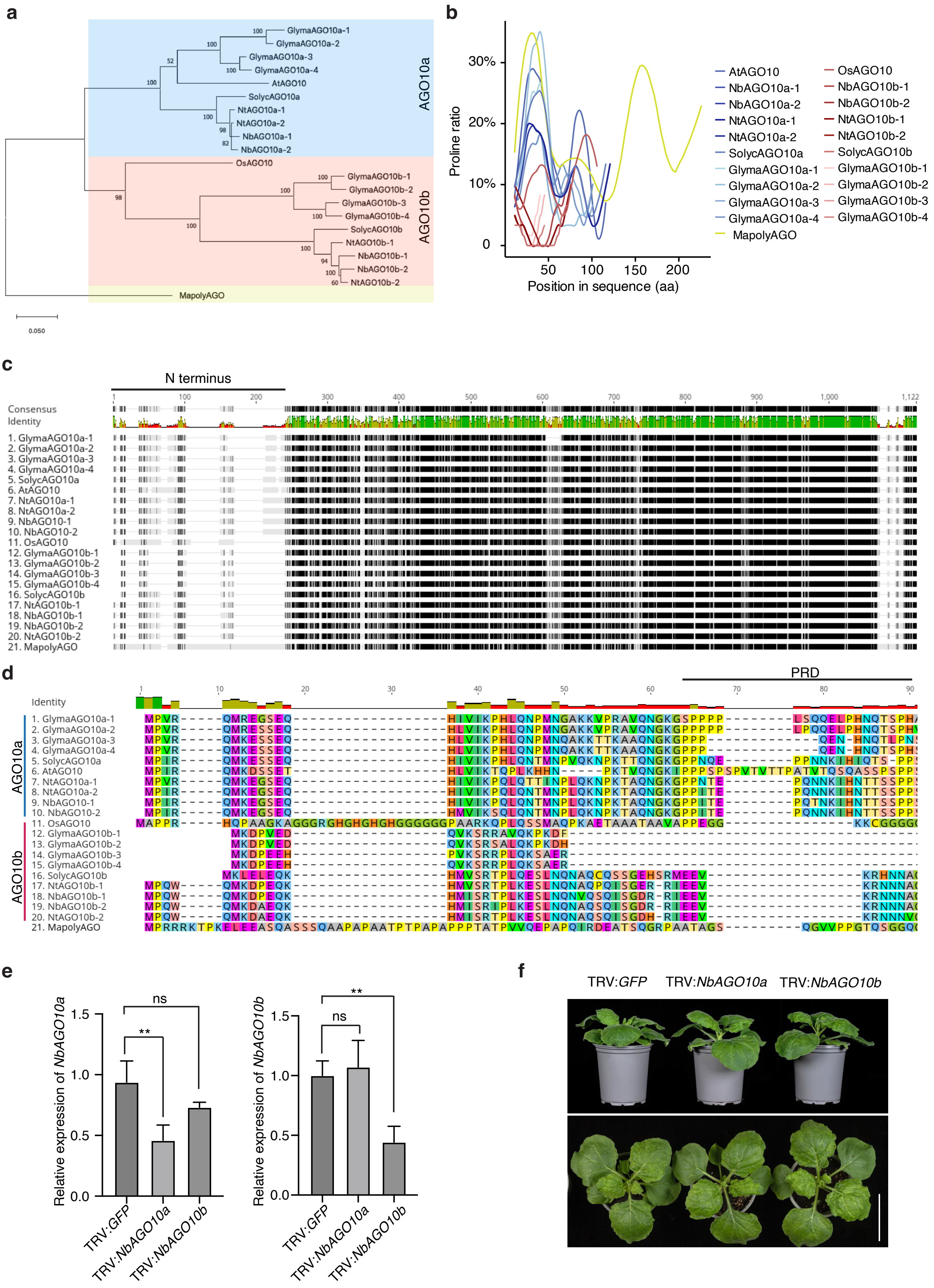

**Supplementary figure 8: A conserved function of AGO10a subclade members in plant immunity.**

**a,** Phylogenetic analysis of AGO10 homologs showing two subclades – AGO10a and AGO10b. AGO10 homologs from representative plant species, including *A. thaliana* (At), *Glycine max* (Glyma), *Marchantia polymorpha* (Mapoly), *N. benthamiana* (Nb), *Nicotiana tabacum* (Nt), *Oryza sativa* (Os), and *Solanum lycopersicum* (Solyc), were analyzed. The liverwort Marchantia encodes one protein in the AGO1/5/10 clade (Mapoly0001s0149 or MapolyAGO in the tree), which was used as the outgroup. The phylogenetic tree was constructed using the neighbour-joining method in MEGA X(42) with 1,000 bootstrap replicates. **b,** Distribution of proline residues in the N-terminal region shows distinct patterns in AGO10a and AGO10b subclades, which are indicated by blue and red, respectively. The yellow line represents MapolyAGO, and the dashed line represents OsAGO10. The proline ratio was calculated using a sliding window of 10 amino acids with a 5-amino-acid step size. **c,** Multiple sequence alignment of AGO10 proteins from different plant species using ClustalW(43) shows a high-level variation in the N-terminal region. **d,** An enlarged view of the N-terminal region highlighting differences in the predicted proline-rich domain (PRD) between AGO10a and AGO10b proteins. **e,** *NbAGO10a* and *NbAGO10b* were efficiently silenced in *N. benthamiana* using VIGS. Transcript abundances were analysed by RT-qPCR with *NbEF1α* as the internal reference. The VIGS vector carrying *GFP* was used as the negative control. Data represent means from three independent experiments ± SEM. Statistical significance was determined using two-tailed Student's t-test (**$p < 0.01$; ns, not significant). **f,** *NbAGO10a*- and *NbAGO10b*-silenced *N. benthamiana* plants exhibited normal growth. Representative images of plants subjected to VIGS targeting *NbAGO10a* or *NbAGO10b*, respectively. Plants infected with the VIGS vector carrying GFP were used as the control. Scale bar, 5.0 cm.
